# “Morphogen gradients applied basally to human embryonic stem cells to control and dissect tissue patterning”

**DOI:** 10.1101/2025.10.30.682158

**Authors:** Tom Wyatt, Mingfeng Qiu, Julie Stoufflet, Hassan Omais, Gabriel Thon, Sara Bonavia, Pascal Hersen, Vincent Hakim, Benoit Sorre

## Abstract

Morphogen gradients are used repeatedly during development to pattern embryonic tissues. Absolute concentrations, duration or even temporal derivative of morphogen concentration have all been proposed to carry positional information. However, establishing the causal relationship between the gradient spatio-temporal profile and resulting cellular diversity and tissue patterning is difficult to address in live embryos because of lack of tools to control those variables.

Here, we developed microfluidics devices able to apply spatio-temporally well-defined morphogen landscapes on the basal side of human embryonic stem cells colonies, thus mimicking how BMP4 is delivered to the pluripotent epiblast during mouse gastrulation. Combining live imaging and theoretical modelling, we show that in this configuration the absolute concentration of BMP4 provides positional information and the cell identities emerging during differentiation can vary according to the shape of the gradient.

Our toolbox provides a powerful means to dissect the logic of patterning and to engineer tissues precisely.

## Introduction

In mammalian embryos, gastrulation is the process during which cells of the pluripotent epiblast are allocated into three germ layers (ectoderm, mesoderm and endoderm) and the body plan is laid out. In the mouse embryo, gastrulation is initiated by BMP4 secreted by the Extra Embryonic Ectoderm (ExE) which creates a concentration gradient on the posterior proximal side of the adjacent epiblast^1^. In response to BMP4 stimulation, epiblast cells start to produce secondary morphogens such as WNT and NODAL, initiating the formation of the Primitive Streak (PS), a structure that extends on the posterior side of the embryo as gastrulation proceeds and from which mesoderm and endoderm emerge, see Fig.1AB. It has been challenging to disentangle the complex interactions between key signalling pathways driving gastrulation and to establish the causality link between their spatio-temporal dynamics and the resulting patterning of the PS^2,3^. This is partly due to the difficulty of observing the embryos at those stages for technical reasons as well as ethical reasons in case of Human, but also to the lack of tools to manipulate signalling landscapes with sufficient spatio-temporal resolution.

**Figure 1:**
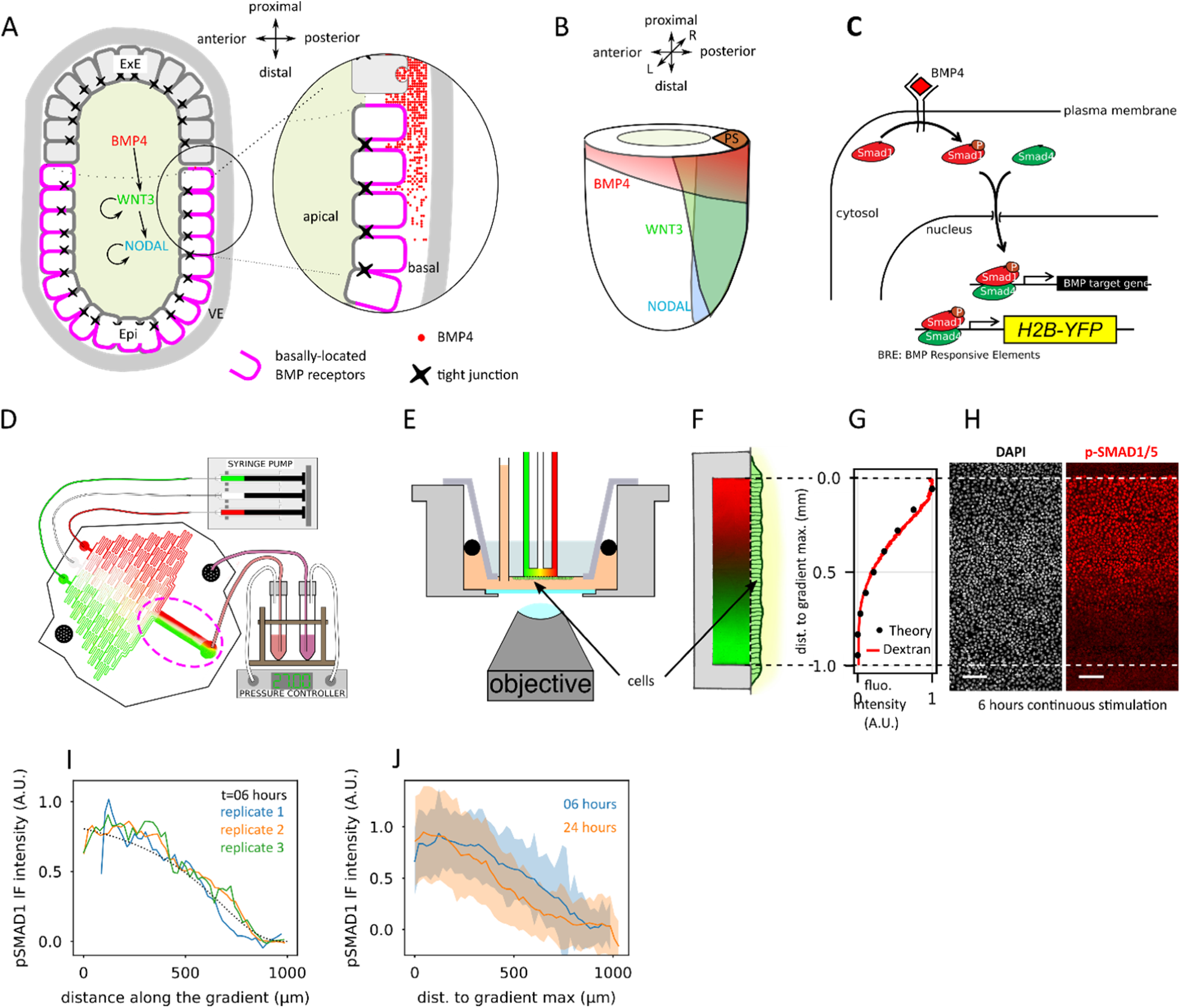
basal stimulation allows to impose a stable gradient of BMP activity on an hESC colony. **A)** Schematic representation of a sagittal section of the mouse embryo at the start of gastrulation (E5.5-E6). BMP4 molecules secreted by the Extra Embryonic Ectoderm (ExE) start a signalling cascade in the adjacent Epiblast (Epi). VE: Visceral Endoderm. Right: close up view revealing the apico-basal polarity of epiblast cells and the restriction of BMP receptors on the basal cells (purple lines). Only the BMP4 molecules present on the basal side of epiblast cells have access to the receptors. They form a proximo-distal gradient in the basal lamina between the epiblast and the surrounding visceral endoderm. Adapted from^9^. **B)** Resulting signalling landscape in the mouse epiblast at mid-gastrulation (E7.0) adapted from^17^. Fate mapping experiments have shown that cells emerging early on from the proximal/posterior region of the PS will migrate proximally to form extra-embryonic mesoderm or laterally to form early embryonic mesoderm derivatives, whereas cells emerging later, i.e. distally to the source of BMP4, will form axial mesoderm and definitive endoderm. **C)** Simplified representation of the BMP signalling cascade. Binding of BMP ligands to their receptor triggers the phosphorylation of SMAD1/5 which then associates with SMAD4 and translocates into the nucleus where the complex binds to DNA through BMP Responsive Elements (BRE) to modulate the expression of its target genes. These BRE have been cloned into a transgenic fluorescent reporter system stably integrated in the genome of hESCs by a PiggyBAC transposase. **D)** General principle diagram of the device. Through successive mixing of 3 media pushed at a constant flow rate in the chip by a syringe pump, a smooth concentration gradient is generated in the rectangular chamber, inside the dotted purple ellipse. A pressure controller sets pressures on the apical and basal sides of the tissue independently to prevent bubble appearance and tissue detachment from the membrane. **E)** Cross section of the gradient chip in its observation cuvette. The morphogen gradient is established in the top compartment while cells are in the bottom compartment. Cells adhere to the porous membrane through which they sense the gradient on their basal side. **F)** Close-up view on the region of the chip where cells are seeded. Rotated by 90° compared to panel E. Parabolic antiparallel gradients in the chip are visualized with green and red fluorescent dextrans (40kDa). **G)** Quantification of the red dextran gradient (red curve) showing good agreement with the theoretical parabolic profile (black dots). **H)** Immuno-fluorescence (IF) of hESC tissue for pSMAD1/5 after 6 hours of stimulation with a parabolic BMP4 gradient ([BMP4]_max_=5ng/ml). **I)** Quantification of pSMAD1 intensity of individual cells as a function of their position in the BMP4 gradient, after 6 hours. Average fluorescence profiles of 3 independent replicates are displayed, showing reproducibility. The dotted line corresponds to the expected pSmad1 profile given a parabolic BMP4 gradient and the measured dose-response curve to BMP4 stimulation of hESCs in sparse cultures: *pSMAD*1 = 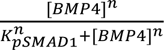 with *K*_*pSMAD*1_ = 0.82*ng*/*ml*, *n* = 1.36. **J)** Comparison of the pSMAD1 profiles at 6 and 24 hours. Solid lines are the average fluorescence of individual cells from 3 and 5 independent experiments at 6 and 24 hours, respectively. Shaded areas represent the standard deviation of the cell populations (n=17×10^3^ and 47 ×10^3^ cells in 3 and 5 replicates respectively) Scale bars: 100µm.

Recently, BMP4 stimulated human Embryonic Stem Cell (hESC) colonies that are confined in 2D on micropatterned substrates and self-organise in concentric rings of the 3 germ layers (h-2D-gastruloids) have emerged as an elegant solution to the accessibility issue^4^. The patterning relies on self-organisation of BMP signalling which is gradually restricted to the edge of the colonies despite homogenous stimulation. This system has permitted to observe and dissect the BMP◊WNT◊ NODAL signalling cascade at play^2,5–7^. However, it does not allow to take control of the spatio-temporal dynamics of BMP signalling in the colony. Microfluidic delivery of an apical gradient of BMP4 over colonies could break the radial symmetry of patterning but didn’t prevent self-organisation^8,9^. One important factor in this self-organisation is the loss of accessibility of the apically provided BMP4 molecules to the BMP receptors located on the basolateral side of the tissue, below tight junctions^10^. In the mouse embryo, for example, ExE secreted BMP4 forms a concentration gradient on the basal side of epiblast cells as it diffuses in the basal lamina between the epiblast and the visceral endoderm, Fig.1AB^9^.

We hypothesised that this configuration could be crucial to define how the morphogen is perceived. Thus, to mimic the *in vivo* situation, we have developed two novel microfluidic devices capable of stimulation of hESC tissue on their basal side with high spatio-temporal resolution. Under static parabolic gradients of BMP4 we have observed a sharp and reproducible pattern of only three cell identities with both fate transitions happening at constant BMP4 concentrations. By contrast, stimulation by a step gradient of BMP4 permitted emergence of the endoderm while this identity was not observed in parabolic gradients. Combining live reporters of signalling activities in the tissue and mathematical modelling, we explain these different behaviours by the dynamics of signalling and especially a bistability in WNT/NODAL signalling, which is strongly constrained by our observations.

## Results

### A microfluidic device for basal stimulation of hESCs with controlled spatio-temporal profiles of morphogens

We designed a two-layer microfluidic chip, in which BMP4 is presented on the cells basal side, through a porous membrane. The top compartment is composed of a microfluidic circuit generating a parabolic gradient of BMP4 concentration in the downstream culture chamber by successive mixing of media inputs, Fig.1D. This parabolic gradient is achieved if one of the side inputs is fed with a medium containing BMP4 while the two others are fed with base medium alone, with equal flow rates. Depending on the configuration of media inputs, linear, dumbbell or even antiparallel gradients of two morphogens can be generated with the same design^11^ see Fig1, Fig.S1, movie 1 and methods.

We verified using fluorescently tagged dextran of a molecular weight comparable to BMP4 (40kDa) that the gradient had a parabolic profile in agreement with the theoretical one (Fig.1FG). As gradient generation does not rely on diffusion, a stable gradient was created in under 1 minute and was stable as long as the flow was maintained, Fig.1D-G & movie 1.

To achieve homogenous and efficient response to BMP4 stimulation, we found that high porosity membranes such as anopore aluminium oxide membranes were necessary, see methods and Fig.S1. As methods commonly used to incorporate membranes in microfluidic devices^12,13^ could not be used with these membranes, we have adapted multilayer soft lithography thermal bonding^14^ to robustly attach the gradient-generating PDMS chip to anopore membranes. To that end, the membranes were first stamped with a thin layer of PDMS to block the pores everywhere except on a rectangular window matching the size of the gradient chamber of the top layer chip^15^. After partial curing, the microfluidic gradient-generating chip was aligned on the stamped membrane and the assembly further cured for irreversible bonding, see Fig.S1A and methods.

A monolayer of hESCs was seeded on the membrane on the opposing side of the gradient window, in the bottom compartment of the device (Fig.S1BC, movie 1 and Methods). hESCs were able to grow and maintain pluripotency in the device for at least 72 hours. We first applied a parabolic gradient of BMP4 ([BMP4]_max_ = 5ng/ml) on the basal side of hESC colonies and assayed cell response by immuno-fluorescence (IF) against phospho-SMAD1, an effector of the pathway immediately downstream of the receptor^16^, Fig.1C. 6 hours after initiation of stimulation, a graded pSMAD1 signal was observed along the BMP4 gradient. Quantification showed that the response profile was reproducible across replicates and followed quantitatively the expected profile given the parabolic BMP4 gradient and the measured dose-response of isolated cells Fig.1HI and S1D. The same quantification after 24 hours of stimulation yielded a similar pSMAD1 profile, Fig.1J. Together, these experiments show that, unlike in apical stimulation experiments where cells’ response to BMP4 self-organises to restrict itself at the edge of the colonies after 24 hours^8–10^, basal gradient stimulation allows one to impose a spatio-temporal profile of BMP stimulation on a colony of hESCs because it preserves accessibility to BMP receptors over time.

### hESCs can interpret absolute concentration of basal BMP4 stimulation

We investigated how the hESC colony would differentiate under a static parabolic gradient with [BMP4]_max_ = 5ng/ml. To reveal the cell identities in real time, we used the Germ Line Reporter (GLR) RUES2 hESC subclone engineered to express fluorescent reporters under the control of genes characteristic of the 3 germ layers: SOX2-mcitrine for pluripotent cells, TBXT-2A-mCerulean for the PS/mesoderm, and SOX17-2A-tomato for the endoderm, hereafter abbreviated as SOX2-YFP, TBXT:CFP and SOX17:RFP^18^, respectively, Fig.2A.

**Figure 2:**
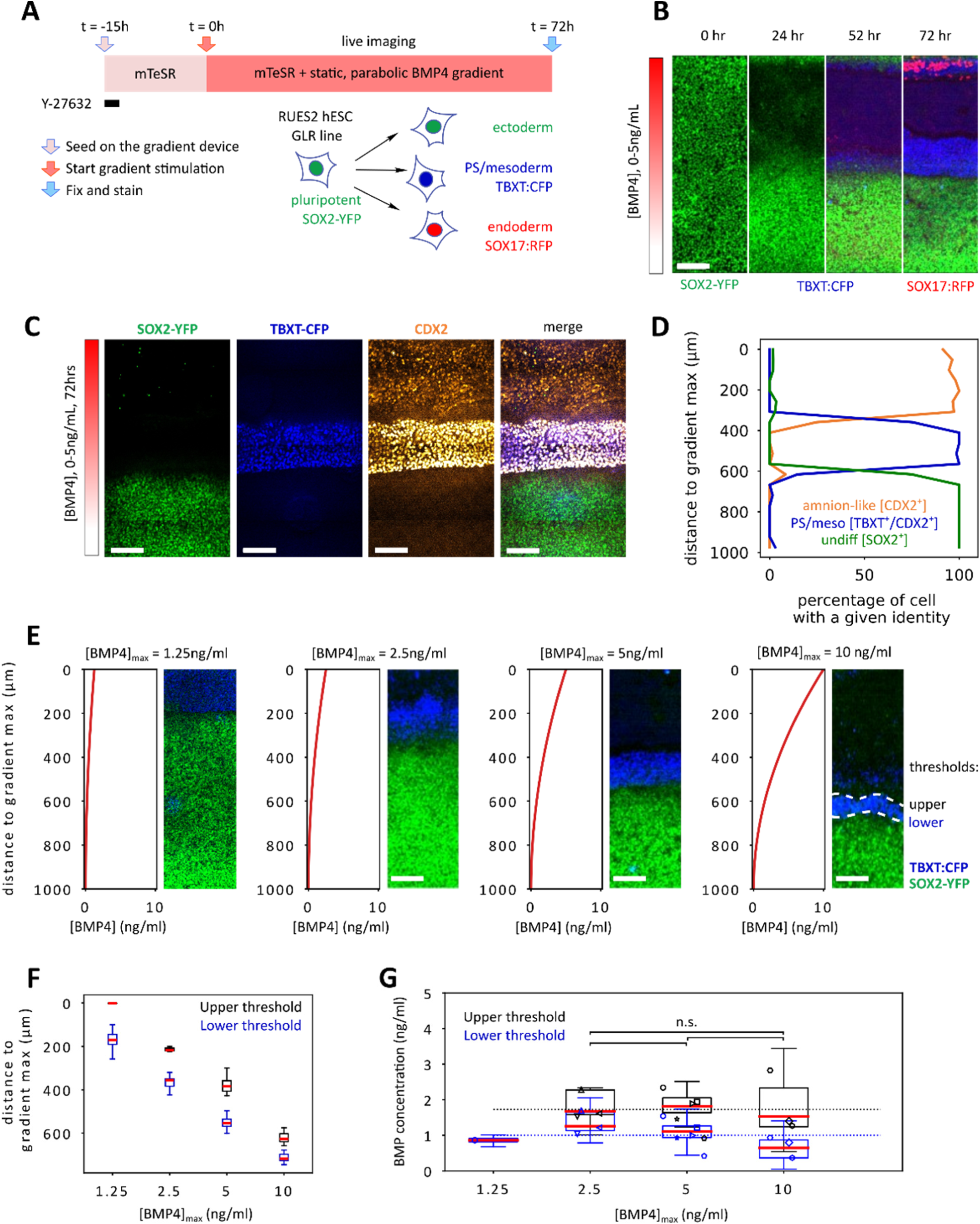
pattern of differentiation of hESCs in a static parabolic gradient of BMP4. **A)** Presentation of the experimental timeline (top left) and of the Germ Layer Reporter (GLR) RUES2 hESC cell line we used in this figure. This cell line has been engineered to report cell identities with the following fluorescent proteins: SOX2-mCitrine (SOX2-YFP) for pluripotency, TBXT-2A-mCerulean (TBXT:CFP) for Primitive Streak (PS)/mesoderm and SOX17-2A-tdTomato (SOX17:RFP) for endoderm. **B)** Temporal evolution of the SOX2-YFP, TBXT:CFP and SOX17:RFP reporters over 72 hours of differentiation in the static parabolic gradient of BMP4 with [BMP4]_max_ = 5ng/ml. **C)** Representative images for the SOX2-YFP, TBXT:CFP and IF for CDX2 after 72 hours in a static parabolic gradient of BMP4 with [BMP4]_max_ = 5ng/ml. **D)** Quantification of the percentage of the cells classified as undifferentiated [SOX2^+^/TBXT^-^/CDX2^-^] PS/mesoderm [SOX2^-^/TBXT^+^/CDX2^+^] and extraembryonic/amnion-like (SOX2^-^/ TBXT^-^/ CDX2^+^) along the gradient for one experiment. **E)** Evolution of the position of the band of TBXT^+^ cells at t=72 hours for 4 values of [BMP4]_max_. The theoretical BMP4 concentration profiles are depicted on the left. **F)** Quantification of evolution of the upper and lower thresholds of the band of TBXT^+^ cells at t=72 hours for 4 values of [BMP4]_max_. The box plots represent the variability of the position of the thresholds. **G)** Evolution of BMP4 concentration at the position of upper and lower thresholds of the band of TBXT^+^ cells as a function of [BMP4]_max_. Red bars represent the median and box plots represent the variability of thresholds position of all replicates for a given [BMP4]_max_. Pairs of black and blue symbols represent the median position of upper and lower thresholds for individual replicates. p-values were calculated using the medians of individual replicates (n=3,5,3 for [BMP4]_max_ = 2.5, 5, 10 ng/ml respectively). For [BMP4]_max_ = 1.25 ng/ml, the upper threshold is not reported as it coincides with the gradient window. Threshold for significance p<0.05. Dotted horizontal lines represent the mean of lower and upper threshold concentrations of all replicates, occurring 1.0±0.1 and 1.7 ±0.2 ng/ml respectively (mean±s.e.m, n=12 independent experiments). Scale bars: 150µm.

Initially all cells were positive for SOX2-YFP, consistent with their pluripotent identity. During the first day of stimulation, SOX2-YFP disappeared at the higher half of the gradient. After 36-42 hours, TBXT:CFP started to be detected in a band of SOX2^-^ cells adjacent to the SOX2^+^ domain (Figs.2B,S2 and movie 2). After 72 hours of stimulation, further characterization with IF revealed that [SOX2^-^/TBXT^-^] cells at the higher end of the gradient were positive for CDX2 indicating an amnion-like cell identity as observed in other BMP4-induced hESC differentiation systems^19–24^. TBXT^+^ cells were also positive for CDX2, indicating a proximal derivative of the PS identity such as extraembryonic mesoderm^25^, also reported after BMP4 stimulation of hESCs and mouse epiblast like cells^26–29^. Characterization of cell identity at the single cell level showed that each domain was composed of an almost pure population of cells with sharp boundaries, Figs.2CD,S2EF.

We then asked how the observed pattern depended on gradient properties. Increasing the maximal concentration while keeping the parabolic shape of the gradient induced a shift of the position of the band of TBXT^+^ cells, Fig.2EF. Converting the position of the boundaries between the domains into BMP4 concentration showed that, within experimental precision, fate transition occurred at a constant concentration, Fig.2G. At [BMP4]_max_ = 1.25ng/ml the [TBXT^-^/CDX2^+^] domain was not induced but TBXT^+^ domain was still present showing that the presence of the [TBXT^-^/CDX2^+^] domain is not necessary to induce the TBXT^+^ cell identity. As in other systems, the emergence of TBXT^+^ cells was dependant on WNT signalling as the band disappeared in the presence of WNT secretion inhibitor IWP2, Fig.S2G.

These experiments show that in static BMP4 gradients, cells can interpret absolute BMP4 concentration to generate 3 domains with sharp boundaries at constant concentration thresholds. The TBXT^+^ cells appeared between 1.0±0.1 and 1.7 ±0.2 ng/ml *i.e.* in the linear range of the dose response of cells to BMP4 (Fig.S1). This value must be compared to the 50ng/ml BMP4 typically used to elicit similar fates with apical stimulation^4,8^. Another important difference with apically stimulated systems is that the repertoire of accessible fates in a static basal gradient is restricted to only 3 identities. Indeed, no SOX17^+^ cells were observed in the static gradient while almost all derivatives of PS observed in h2D-Gastuloids^30^. SOX17^+^ cells, mixed with TBXT^+^ cells, were nevertheless observed, but outside the stimulation area at the edge between the high end of the gradient and the unstimulated zone, where the tissue experiences a step gradient of BMP4 concentration Figs.2B,S2AB & movie S2.

### An Endoderm tube sheathed with mesoderm cells emerges at the edge of BMP4 stimulated area

To dissect what happens at the edge of the stimulation window, we have developed a simplified variant of the gradient chip, called µTransWells, creating patches of homogenously basally stimulated cells in an hESC colony. Like the basal gradient chip, this device is fabricated by stamping PDMS on an anopore membrane to block its pores everywhere except in 700µm-diameter disk-shaped windows, on top of which 4mm-diameter PDMS wells are attached for individual stimulation, Fig.3AB. IF against pSMAD1 after 24 hours of BMP4 stimulation confirmed that this device was able to produce a patch of homogeneous, BMP4 stimulated cells with a sharp boundary, Fig.3CD.

**Figure 3.**
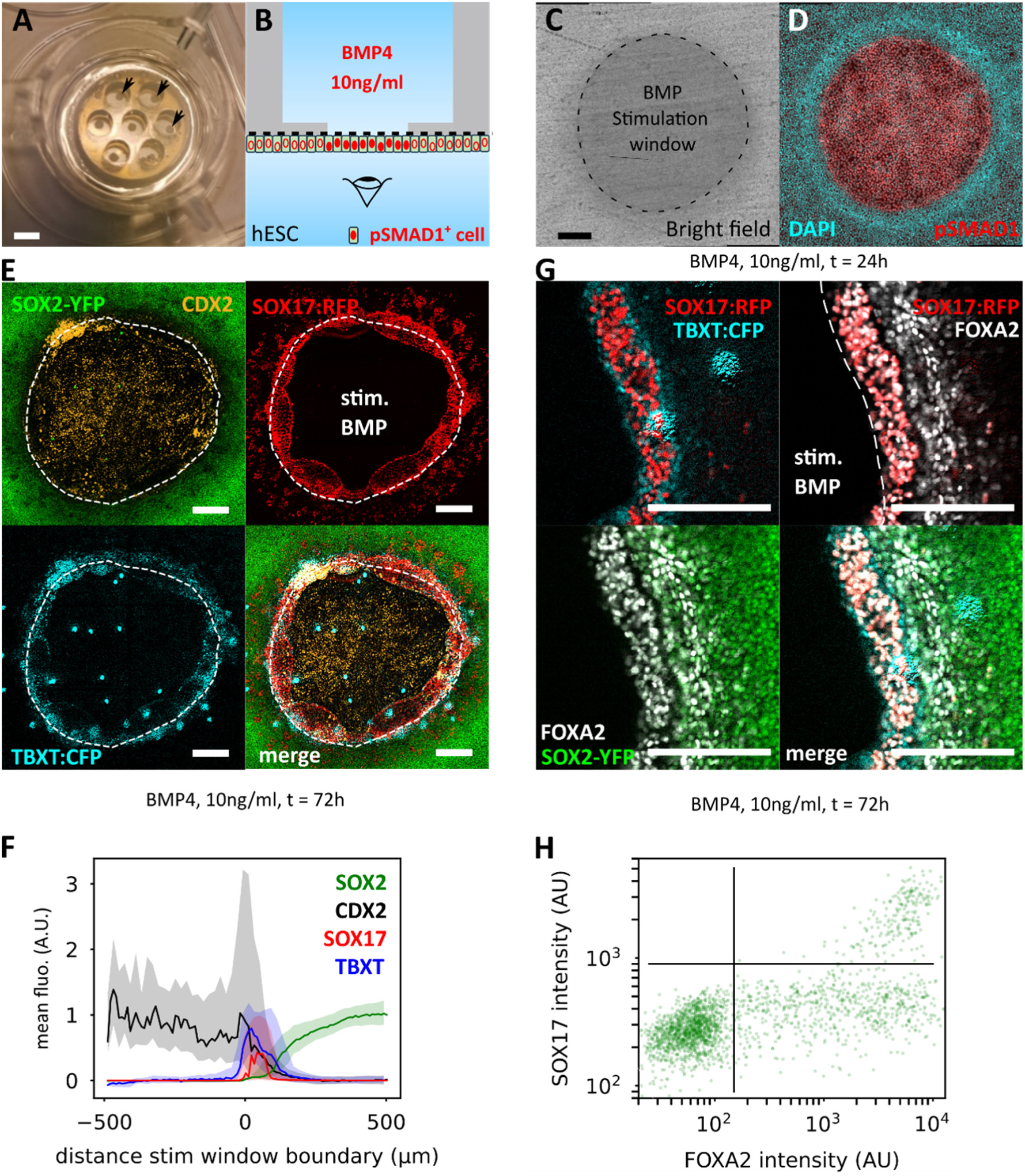
Production of well-defined signalling centers on hESC epithelia with the µTransWell device. **A)** Top view of a µTransWell device, with 4 mm-diameter small wells, each with a 700µm-diameter circular stimulation patch (arrows). Scale bar 4mm. **B)** Cross section of the culture system showing the experimental configuration. Cells are attached to one side of the membrane in the bottom compartment and BMP4 is provided in the top compartment through the membrane but only the cells in the area where the membrane is not blocked by PDMS receive the morphogen (pSMAD1 positive cells, depicted with red-filled nuclei). Not to scale. Microscopy observation is achieved from the bottom. **C,D)** Image of one µTransWell stimulation patch. **C)** Bright field. The darker-grey area is the non-obstructed part of the membrane, where cells can receive BMP4. **D)** immunofluorescence against pSMAD1 after 24hours of BMP4 stimulation on the basal side of the central patch in the tissue. Scale bar 150µm. **E)** Representative fluorescence images for the SOX2-YFP, TBXT:CFP and SOX17:RFP GLR reporter line, plus immunofluorescence for CDX2 after 72h of stimulation of a central patch of the tissue with [BMP4] = 10ng/ml. Scale bar 150µm. **F)** Average fluorescence profiles after 72hours of BMP4 stimulation as a function of the distance to the edge of the stimulation window. Colored areas represent the first and third quartile of individual cells from 3 independent replicates. **G)** Close-up view of the ring of differentiated cells at the edge of the BMP4 stimulation area after 72hours of BMP4 stimulation plus immunofluorescence for FOXA2. SOX17:RFP^+^ cells co-express FOXA2 and are surrounded by TBXT:CFP^+^ cells. scale bar 150µm. **H)** Quantification of FOXA2 and SOX17 co-expression after 72hours of BMP4 stimulation. Each dot represents a single cell. All SOX17:RFP^+^cells are also positive for FOXA2 (upper right quadrant).

The device robustly reproduced the phenotype observed at the top edge of the gradient chip, *i.e.* after 72 hours, cells on the stimulation window were [CDX2^+^/ISL1^+^/GATA3^+^/TFAP2A^+^] confirming the amnion-like identity of cells receiving a high dose of BMP4 (10ng/ml), unstimulated cells remained [SOX2^+^/OCT4^+^] and in between emerged a thick domain with TBXT^+^ and SOX17^+^ cells, Fig.3EF,S3. Observation at higher magnification revealed that this domain had a reproducible 3D structure with a hollow tube of SOX17^+^ cells surrounded by TBXT^+^ cells. SOX17^+^ cells co-expressed FOAX2, confirming their endoderm identity Fig.3GH, movie 3.

TBXT^+^ cells were detected on the stimulation window for lowered BMP4 concentrations between 0.6 to 2.5 ng/ml *i.e.* in a range comparable, although a bit wider, to the concentration thresholds in the gradient chip (Figs.2, S3AB). However, unlike in the gradient chip, TBXT^+^ cells were always observed as patches, in variable amount, never covering more than 60% of the stimulation window, Fig.S3AB. By contrast, in WNT3a stimulated µTransWells colonies, 100% of the cells on the stimulation window were TBXT^+^. In that condition, SOX17^+^ cells also appeared at the frontier of stimulated and unstimulated domains but as individual motile cells rather than as a close packed tube, figS3C, movie 4.

### Transient BMP4 stimulation triggers sustained WNT and NODAL signalling and permits endoderm specification

We hypothesised that the emergence of TBXT^+^ and SOX17^+^ cells at the edge of the stimulation windows was due to endogenous WNT and NODAL signalling as it has been reported in the embryo and in other *in vitro* peri-gastrulation systems that BMP4 stimulation induces production of WNT3 and NODAL ligands^7,31,32^. Indeed, the band of TBXT^+^ and SOX17^+^ cells disappeared upon treatment with the WNT secretion inhibitor IWP2, Fig.S3F.

We then investigated the spatio-temporal dynamics of activation of the BMP, NODAL and WNT signalling pathways using IF and signalling reporter cell lines. As before, pSMAD1 stain was strictly restricted to the stimulation area after 24 hours of BMP4 stimulation while cells positive for BRE:YFP, a transgenic reporter of BMP activity^33^ (Fig.1C&S4A), could be detected outside the stimulation area, Fig.4AB. The apparent discrepancy between the BRE:YFP and pSMAD1 signals can easily be explained by their relative lifetime. Indeed, inhibition of BMP signalling with LDN-193189 showed that pSMAD1 IF was back to baseline 30 minutes after inhibition while BRE:YFP was only reduced by 15% compared to control after 24 hours (Fig.S4BC). This suggests that the BRE:YFP^+^ cells outside the stimulation have received BMP4 in their past. Time lapse imaging clearly showed that cells move outside the stimulation area, movie 5. Single cell tracking experiment confirmed this hypothesis: 100% of tracks of BRE:YFP^+^ cells found outside the stimulation window after 24 hours of stimulation originated from the stimulation area, on which they had spent on average 11+/-5 hours, Fig4.CD.

**Figure 4:**
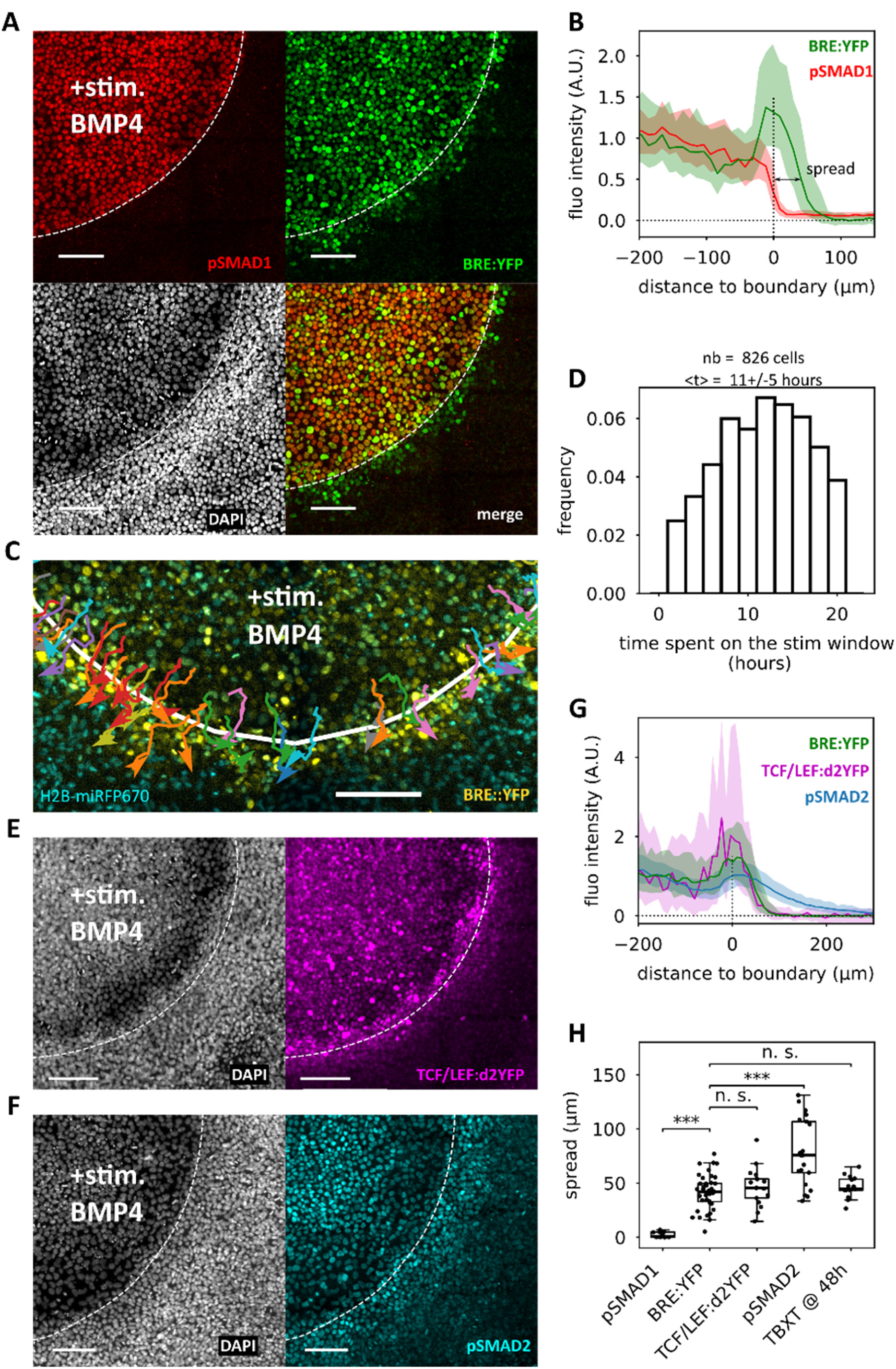
Differentiated cells outside the stimulation window have received a pulse of BMP4 that has activated WNT and NODAL signalling. **A)** Comparison of pSMAD1 immunofluorescence stain and BRE:YFP signal after 24 hours of stimulation of the central patch of cells with BMP4 (10ng/ml). The white dotted line represents the boundary between stimulated and non-stimulated regions of the tissue. **B)** Average BRE:YFP and pSMAD1 fluorescence profiles after 24 hours of stimulation of the central patch of cells with BMP4 (10ng/ml), as a function of the distance to the edge of the stimulation window. Fluorescence profiles are normalized by the average of the signal within the stimulation window. Colored areas around the curves represent the first and third quartile of individual cells. From these plots the spread of signalling is extracted, defined as the distance at which the signalling curve becomes smaller than 0.5. **C)** Single cell trajectories of BRE:YFP^+^ cells and outside the stimulation window after 22hours of stimulation, showing that all cells fitting both criteria have spent time in the stimulation window and thus have received BMP4. **D)** Histogram of time spent on the stimulation window by BRE:YFP^+^ cells positive and outside the stimulation window 22hours after the beginning of stimulation. N=826 cells from 12 colonies. **E,F)** TCF/LEF:d2YFP (E) and pSMAD2 (F) Immunofluorescence stain after 24 hours stimulation of the central patch of cells with BMP4 (10ng/ml). **G)** Average BRE:YFP, TCF/LEF:d2YFP and pSMAD2 fluorescence profiles extracted from a single colony after 24 hours of stimulation of the central patch of cells with BMP4 (10ng/ml), as a function of the distance to the edge of the stimulation window. Colored areas represent the first and third quartile of individual cells from 3 independent replicates. **H)** Box plots representing the measured spread of signalling outside the stimulation area (as defined in B). Dots represent individual replicates. Signalling reporters spread (pSMAD1, BRE:YFP, pSMAD2, TCF:d2YFP) have been measured after 24 hours of BMP4 stimulation while position of the max of TBXT stain was assayed after 48 hours. Threshold for significance: p-value of student’s t-test <0.05. Scale bars 100µm.

To record the activity of the WNT/β-catenin pathway with a higher temporal resolution than the BRE:YFP system, we engineered a transgenic reporter cell line expressing a destabilized YFP (d2YFP) under the control of the TCF/LEF synthetic reporter^34^ (TCF/LEF:d2YFP). After 24hours of BMP4 stimulation, TCF/LEF:d2YFP signal was elevated on the BMP4 stimulated area and spread about 50µm outside. By mixing BRE:YFP and TCF/LEF:d2FFP cells in the same experiment (BRE:YFP cells having a Far Red nuclear marker to distinguish them) we observed a strong correlation between the BRE and TCF spread at the single colony level, Fig.S4J. ACTIVIN/NODAL pathway activity was assayed by IF against pSMAD2. It was also found to be high on the stimulation window and spread further from its boundary than the BRE:YFP, TCF:d2YFP or TBXT signals (∼70µm), Fig4FGH&S4.

At 48 hours, the ring of TBXT^+^ cells was observed within the spread of BRE:YFP signal, showing that these cells have experienced BMP4 stimulation in their past, FigS4K-O. Taken together, these observations suggest that emergence of TBXT^+^ cells outside the stimulation window results from sustained WNT and NODAL triggered by transient exposure to BMP4 and that WNT is the limiting factor for the differentiation of TBXT^+^ cells. This condition permits the emergence of SOX17^+^ definitive endoderm cells while the static gradient does not.

### Patterning logic is well captured by a simple cell fate network with bistable WNT/NODAL signalling

We then asked if the logic of patterning observed could be consistent with the reported effect of BMP4 stimulation in the mouse embryo and in other peri-gastrulation *in vitro* systems^24^. We thus designed a minimal Cell Fate Network incorporating the following interactions: BMP4 induces directly amnion-like cells and the production of WNT and NODAL ligands^7,31,35^ which promote induction of endoderm and mesoderm cells^36^. Amnion-like and mesoderm identities cross-repress each other, and BMP4 prevents the emergence of endoderm cells^28^. WNT and NODAL signalling self-activate^7,24,31,35^. The fact that TBXT is found only in the area positive for both TCF/LEF:d2YFP and pSMAD2, suggests that WNT is limiting for differentiation, therefore as a simplification, the model considers the 2 morphogens as a single entity represented by WNT, Fig.5B and mathematical supplement. Considering their individual contribution would be necessary, however, to resolve the fate choice between endoderm and mesoderm.

**Figure 5:**
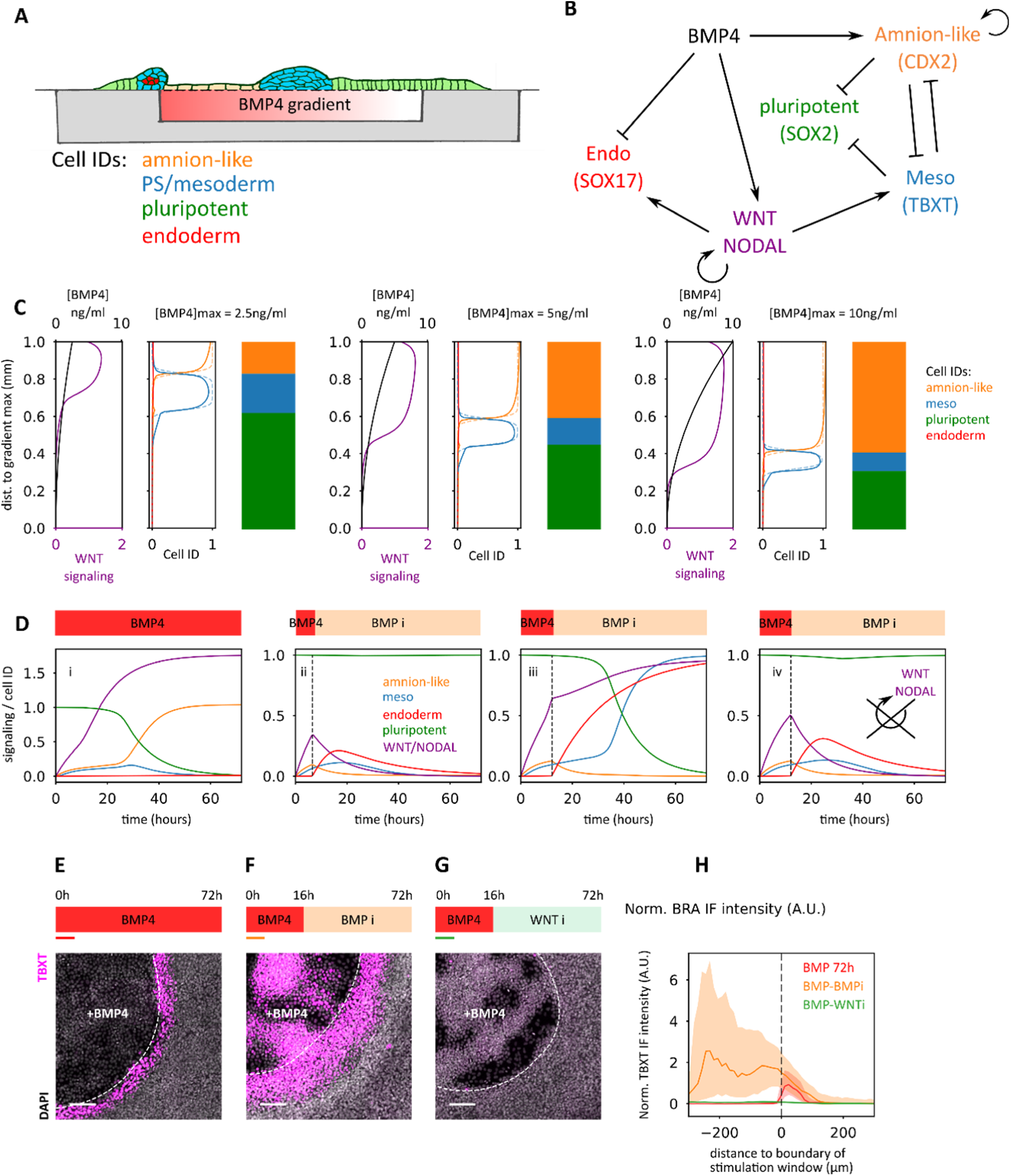
Model for patterning – Bistability of WNT. **A)** Schematic summary of our observations. In the static gradient only 3 cell identities are observed. From minimum to maximum BMP4 concentration: undifferentiated, PS/mesoderm and amnion-like cells. Outside the stimulation area, at the edge between un-stimulated cells and those experiencing the high end of the gradient: mesoderm and endoderm are specified, dueto WNT and NODAL endogenous signalling triggered by transient BMP4 stimulation. **B)** Proposed Cell Fate Network to explain the logic of patterning in our systems: BMP4 induces extra-embryonic amnion-like cell identity directly but also WNT and NODAL ligands that can induce mesoderm and endoderm fates. However, endoderm is blocked by BMP4. Amnion-like and PS/mesodermal fates cross inhibit. WNT and NODAL are secreted ligands so they can diffuse. WNT and Nodal self-activate. We have included the self-activation of amnion-like cells, which is found to be unimportant through parameter fitting. **C)** Model prediction of patterning in a static gradient. For each [BMP4]_max_ the left panel represents the applied BMP4 concentration gradient and the resulting WNT signalling profile, and the center panel represents the predicted cell fate patterning (solid lines). The right panel represents the expected pattern of cell identities based on experimental observations of figure 2, also denoted by sigmoidal profiles in the center panel (dashed lines). **D)** Model prediction of the outcome of transient BMP4 stimulation as experienced by cells in µTransWell experiments. (i) Cells staying on the stimulation window experience sustained BMP4 stimulation and amnion-like fate prevails. (ii) Transient but shorter than characteristic time to induce WNT self-activation results in no differentiation. (iii) At the edge of the stimulation window, cells experience transient BMP4 stimulation inducing self-sustained WNT signalling and PS/mesoderm identity. (iv) Without WNT self-activation, transient stimulation does not induce differentiation. **E,F,G)** TBXT stains on a µTransWell after 72h BMP4 (CTRL), 16h BMP4 followed by 56h BMP inhibition (BMPi) and 16h BMP4 followed by 56h WNT inhibition (WNTi). After BMPi, TBXT is everywhere showing that sustained BMP4 is needed to get the amnion-like fate. WNTi leads to no TBXT+ cells, indicating that this fate needs sustained WNT activity and therefore that bistable WNT is necessary. Scale bars 100µm. **H)** Average fluorescence profiles for the 3 conditions described in panels EFG for n = (13, 13, 15) colonies respectively, from 3 independent experiments. Colored areas represent the spread between the 1^st^ and 3^rd^ quartiles of the distribution of individual cells. Profiles are normalized to the maximum of the average TBXT profile in the BMP72h condition.

The mutual repression between Amnion-like and mesoderm identities was sufficient to obtain models reproducing, after parameter optimisation, the emergence of 3 domains with boundary at constant concentration thresholds in a static gradient of BMP4, Fig.5C, in a manner similar to the French Flag model^32^, with some significant differences, as described below. To recapitulate the presence of mesoderm and endoderm cells outside the simulation window, the self-activation of WNT/NODAL was crucial to create a bistable behaviour permitting the persistence of WNT and NODAL signalling after BMP stimulation had stopped, Fig.5D, as further detailed in the Mathematical supplement.

The model should also account for the µTransWell experiment data as well as for the time development of the patterns in static BMP4 gradients (Fig.S2). The differentiation dependence upon the duration of BMP4 stimulation in the µTransWell experiments determined the time scale of WNT dynamics to be about δ ^-1^ ∼11 hours (Figs.4D, S4I&S5A). The WNT spatio-temporal dynamics is further constrained by the small extent of the TXBT^+^ region close to the high BMP4 end outside the gradient. This shows that the diffusion length of WNT, *l*_w_=(η_w_/ δ_W_)^1/2^ should be smaller than 50μm and thus that WNT diffusion constant η_w_ should be smaller than 0.06μm^2^/s, in agreement with reported values in other settings^37^.

The bistable behaviour of WNT dynamics, in combination with the parameter constraints, leads to two important differences of our model with the French flag picture. Firstly, the low BMP4 boundary of the TBXT^+^ band is determined by the location of the front between high and low WNT concentrations rather than directly by BMP4 concentration. Secondly, this WNT front moves at a slow speed. The above constraints on WNT dynamics and diffusion were found compatible with the time development of the TBXT fluorescence (Fig.S2) only if the TBXT^+^ band after 72 hrs observed in our experiment (Fig.2) was still very slowly growing at its low BMP4 boundary rather than fully stationary, as in the French flag picture. These arguments are detailed further in the mathematical supplement.

To verify that WNT bistability was operating in our system, we performed inhibition experiments. We observed that TCF/LEF signal persisted after inhibition of BMP signalling with a pharmalogical inhibitor, Fig.S4GI. Inhibition of BMP or WNT signalling had also drastic consequence at the level of cell fate choice. In µTransWells, the ring of TBXT^+^ cells disappeared upon WNT signalling inhibition after 16 hours of BMP4 stimulation. This confirms that persistent WNT activity even when cells have moved outside the stimulation area is necessary for mesoderm specification of those cells, Fig.5GH,S5BC. When BMP signalling was inhibited after 16 or 24 hours of BMP4 stimulation, the amnion-like cells present on the stimulation area in the control were mostly replaced by TBXT^+^ cells. Sustained BMP4 stimulation is thus needed for amnion-like cells specification which in turn prevents mesoderm, Figs.5E-H,S5A.

## Discussion

BMP has been shown to work as a morphogen, *i.e.* controlling cell fate in a dose dependent manner, in a variety of developmental contexts^16^. The situation is less clear for *in vitro* hESC directed differentiation^20,26,38^. Isolated or low-density hESCs can only differentiate in amnion like cells if stimulated with a high enough dose of BMP4 while in denser 2D cultures, mesoderm cells can appear in an intermediate range of BMP4 concentration, but never as a pure population^39^. Finally, cells from the three germ layers are generated in BMP4 stimulated micropatterned hESC colonies^30^. Together, these observations lead to the conclusion that hESCs could not reliably interpret concentration as a predictive differentiation cue^24^.

Here we introduce microfluidic devices that, taking inspiration from where the BMP4 molecules are delivered to the mouse epiblast, allow stimulating epithelial tissues on their basal side with defined spatio-temporal profiles of morphogen. In static parabolic gradients of BMP4, hESCs differentiate into a sharp and reproducible pattern of 3 cell identities with a central band containing 100% of TBXT^+^ mesoderm cells. The two cell fate thresholds happen at constant concentrations, in the dynamic range of the sensitivity of the BMP pathway, thus establishing that absolute concentration of BMP4 can provide positional information for hESCs to solve Wolpert’s French Flag problem^40,41^. Basal stimulation with a spatial gradient, however, is necessary to reveal the morphogen effect as the BMP induced mesoderm domain is not stable over large areas as shown in µTransWells dose response experiments.

Under static gradient conditions however, cells have access to only 2 differentiated identities, amnion-like and extra-embryonic mesoderm. Unlike in h2D-gastruloids^4,30^, other embryonic derivatives of the PS such as definitive endoderm are absent. Gastrulation is a dynamic process during which cells are certainly experiencing dynamic stimulation. Our observation of cell differentiation at the edge between stimulated and unstimulated regions of tissues illustrates possible consequences. Using live reporter of signalling and tracking of cell position, we established that this differentiation was due to cell moving away from the BMP stimulation area, thus experiencing a transient exposure to BMP. Inhibition experiments and modelling further showed that, in line with a recent report^24^, the key regulatory network feature to explain this effect is the bistable expression of secondary morphogens WNT and NODAL.

Thus, besides generating a temporal morphogen effect, modulating the duration of BMP exposure also increased the repertoire of accessible fates, as endoderm cells emerged among those cells transiently exposed to BMP. Prolonged exposure to BMP does not induce endoderm *in vitro*^42^ as this prevents its emergence^28,29^. Short stimulation, however, has been used to induce posterior foregut organoids^43^. Generation of endoderm after a short pulse of BMP is consistent with fate mapping experiments in the gastrulating mouse embryo reporting that endoderm cells can emerge from rather proximal regions of the PS^44^. These cells could experience transient exposure to BMP by migrating away from the source and expressing BMP inhibitors such as CER1. Effectively transient exposure to BMP could also explain the emergence of endoderm cells in BMP stimulated hESC micropatterned colonies^4,30^.

Thus, our basal stimulation devices circumvent a major source of variability in interpretation of BMP concentration, namely the loss of accessibility to receptors of apically provided ligands due to gradual segregation of receptors on the basal side during maturation of the epithelium. They allowed us, combined with live reporters of signalling and cell fates, to establish the causality link between the morphogen landscape experienced by the tissue and the resulting patterning. As morphogen gradients and morphogen-induced signalling centres are ubiquitous in development^45^, and given that in a number of situations such as neural tube patterning, morphogens are provided on the basal side of epithelia, we expect our experimental strategy to be widely applicable to dissect the mechanism of patterning of other developing organs and for precise tissue engineering.

## Supporting information

mathematical supplement

## Methods

### Design of microfluidic device

among the many gradient-making devices available in the microfluidics toolbox (reviewed in^46^) we have favoured a design using successive mixing as it allows for more flexibility in the gradient shape and temporal variation of the gradient. We have thus adapted an original design of Jeon *et al.* ^11^ according to estimated diffusive properties of BMP4. Previous measurements of the diffusion coefficient of freely diffusing BMP and other proteins of comparable molecular weights give an order of magnitude estimate for a diffusion coefficient of *D*_B_ ≈ 50 *µm*^2^*s*^-1 47,48^. This diffusion coefficient constrains the device design due to the requirement that complete mixing by diffusion must occur in each serpentine channel. We thus chose channel dimensions and a flow rate which satisfy the condition that the root-mean-squared distance travelled by BMP4 during passage in a single serpentine channel is greater than half the channel width. That is, that

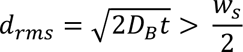

where *t* is the passage time for liquid travelling through a serpentine channel and *w*_s_is the width of the serpentine channel. Dimensions of 70 *µm* ∗ 8300 *µm* ∗ 150 *µm* (width ∗ length ∗ height) and a flow rate (for each of the three inlet channels) of 10 *µL*. *s*^-1^ were thus chosen to satisfy this condition. The chosen flow rate, within the stimulation chamber of dimensions 1000 *µm* ∗ 4000 *µm* ∗ 150 *µm* (width ∗ length ∗ height) results in an estimated root-mean-squared displacement of ∼85 *µm* for BMP during passage through the stimulation chamber. We expect only minimal change in the profile of the gradient from inlet to outlet, since this is small compared with the chamber width.

### Manufacture of PDMS microfluidic chips

The microfluidic design was drawn in vector graphics software Inkscape and imaged onto polyester film with a photographic emulsion coating to create a mask (JD Photo Data). The resin mould for casting the microfluidic chip was created using standard protocols from photolithography. Briefly, SU8 photoresist **(microresist,GmBH)** was spun onto a Si wafer to a thickness of 150 μm and then baked for 1hr at 95°C. The mask was put in contact with the baked resin and exposed to UV for 60 s. This was then baked again for 1hr at 95°C before developing in SU8 developer for 20 min. The mould was then silanized for 1 hr under vacuum using (3-Mercaptopropyl)trimethoxysilane (Sigma). The PDMS microfluidic chip was carefully cut out of the mould using a scalpel and holes were punched in the inlets and outlets.

### Patterned stamp-and-stick to bond the chip to the membrane

Basal delivery of BMP4 to hESC colonies imposes to bind the microfluidic chip to a porous membrane. This is a common feature of organ on chip devices, however in most cases they incorporate PDMS or track-etch membranes with low pore density (10^6^ pores / cm^2^, i.e ∼1 pore./cell on average) which we found were not suitable for our application as BMP stimulation though these membranes resulted in a spatially heterogenous response. We thus used anopore membranes (Whatman anodisc, 0.2µm pores) which have around 80% porosity. As common bonding techniques such as plasma bonding do not work to robustly bond the microfluidic chip to anopre membrane, we combined the stamp-and-stick ^49,50^ and multilayer soft lithography^14^ techniques. Briefly, a thin layer of PDMS was stamped onto the membrane using a custom-made PDMS stamp. The stamp was made by casting PDMS from an SU-8 mould consisting of a single raised rectangle which matched the dimensions of the stimulation window of the gradient chip. Thus, the resulting stamp deposited PDMS onto the membrane everywhere apart from the stimulation window. To ink the stamp, a thin layer of PDMS was spun onto a glass coverslip and the stamp was placed face-down onto the coated glass. The stamp was removed and left in a dust-free environment for 10 minutes to allow inhomogeneities in the PDMS thickness to relax. The stamp was then carefully placed on the membrane, left for 30 seconds to ensure uniform contact, and then carefully removed. The microfluidic chip was aligned by hand under a magnifying glass and pressed against the membrane and backed at 65°C for 1 hr. The stamping of PMDS thus served the dual-purpose of bonding the chip to the membrane and blocking membrane pores in all regions where apical to basal communication should not occur, see Fig S1.

### Microfluidic experiment preparation

PTFE tubing was rinsed with deionised water and then autoclaved. GasTight glass syringes (Hamilton; 0.5 mL) and pressure vials were sterilised in a 70% ethanol bath followed by rinsing in deionised water. The tubing and syringes were filled with the required stimulation media and connected, taking care not to trap bubbles. The microfluidic device was the treated for 7 min with deep UV (Jelight UVO cleaner) and a 70 μL droplet of Laminin-521 (Stemcell Technologies), diluted in PBS+/+ (Sigma) to 20 μg m*L*^-1^, was placed on the membrane, covering the stimulation window, for 45 min, see Fig.S1.

### hPSC cell lines, culture and differentiation

The RUES2 and RUES2 GLR hESC cell lines were kindly provided by A. H. Brivanlou (Rockefeller University, NY, USA). All experiments were declared to the French Agence de la Biomedecine, authorisation number RE16-004RCI and RE21-001RC. hESC were cultured on culture dishes coated in Geltrex (Thermo Fisher Scientific) in mTeSR according to the manufacturer’s guidelines. iPSC029 cell line is a kind gift of Mickael Kyba (University of Minnesota, USA). Cultures were tested for mycoplasma infection once a month. Gradient chip experiments were performed with the RUES2 parental line and the GLR and signalling reporter subclones. µTransWell experiments were performed with the various RUES2 subclones and the iPSC0029 cell line with similar outcome. hESC and hiPSC were cultivated in mTeSR plus medium according to manufacturer’s instruction on tissue culture dishes coated with 1%Geltrex solutions. Prior to seeding on microfluidic devices, hESC were harvested as a single cell suspension using accutase (sigma) incubated for 6 minutes at 37°C a drop of cell suspension (2-3.5.10^6^ cells/ml) in mTeSR medium supplemented with ROCK inhibitor (RI,20µM) was positioned on top of stimulation window of the devices and cells were allowed to attach to the substrate for 45 minutes. The devices were then transferred in the imaging cuvette filled with mTeSR+RI for 45 additional minutes after which the medium was replaced by mTeSR. Stimulation started between 4 to 15 hours after seeding with BMP4 [R&D System, 314-BP], WNT3A [R&D System, 5036-WN] at concentration specified in each experiment. Pharmacological agonists or inhibitors were used at the following concentration unless otherwise specified: CHIR99021 [3µM, cell guidance systems, SM13], LDN193189 [100nM Cell Guidance System SM23], IWP2 [2µM, Stemcell Technologies, 72122], SB431542 [20µM, cell guidance systems, SM33], XAV939[5µM, cell guidance systems, SM38]

### Reporter cell lines

the RUES2 GLR cell line generation and validation has been described in ^18^. It is engineered to report for the expression of three independent fate reporters: SOX2–mCitrine (SOX2-YFP), BRA–2A-mCerulean (TBXT:CFP) and SOX17-2A–tdTomato (SOX17:RFP). Fluorescent reporter proteins were inserted at the endogenous loci using crispr/cas9 gene editing. The BRE:YFP and TCF/LEF:d2YFP clonal and stable transgenic RUES2 lines were generated using the ePiggyBAC transposable elements system ^51^. the BRE element (addgene plasmid#45126, ^33^) was cloned in a ePiggyBAC plasmid containing a puromycin resistance cassette vector (gift of AH Brivanlou) upstream of a minP minimal promoter (from Promega pGL4) and a YFP. The TCF/LEF (addgene #32610 ^34^) was cloned upstream of a hsp68 minimal promoter and d2YFP, a destabilized version of YFP with a lifetime of 2h. Cells were transfected using lipofectamine 3000 following manufacturer’s instruction. Selection with puromycin (0.5µg/ml) was started 48h after transfection. After a week of selection cells were seeded at clonal density (100-200 cells in a 35mm dish) in mTeSR supplemented with CloneR2 (stemcells technologies). Clones were picked manually and assayed for their capacity to respond to BMP4 or CHIR treatment.

### Immunofluorescence

Tissues attached to membranes were fixed by placing the transwell in a 6-well dish containing in 4% PFA for 25 min and were then washed 3 times for 15 min in PBS. They were then blocked and permeabilised in 2% bovine serum albumin with 1% triton-100 (Sigma) for 30 min. The tissue was washed again in PBS incubated with primary antibody in blocking solution overnight at 4°C. The tissue was then washed 3 times for 30 min in PBS before incubating with the secondary antibody for 2 hr at room temperature. The tissue was then washed in PBS and kept in PBS at 4°C until imaging. See table below for a list of used antibodies.

### Imaging

All imaging was performed on an inverted Olympus IX83 equipped with DSD2 confocal module (Andor). Montage scans were performed using a motorised stage (Märzhäuser). Sample was maintained at 37°C and 5%CO2.

### Image analysis

Image processing was performed using custom-made routines. Fluorescence illuminations inhomogeneities were corrected using the BaSIC algorithm^52^, tiles were assembled using the grid/collection stitching Fiji pluggin^53^. Nuclei segmentation was performed with stardist ^54^ and tracking with trackmate ^55^

### Mathematical modelling

The cell fate network model (Fig.5B) is described by the following nondimensional differential equations

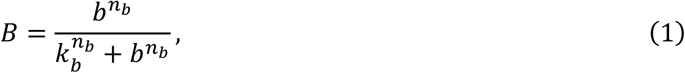

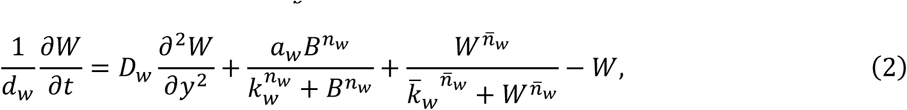

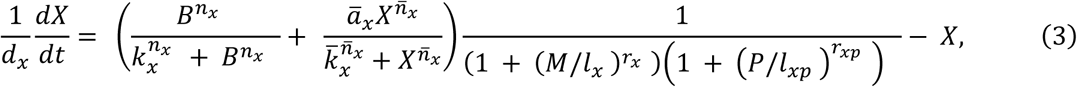

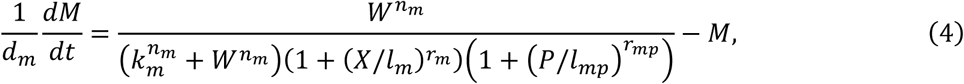

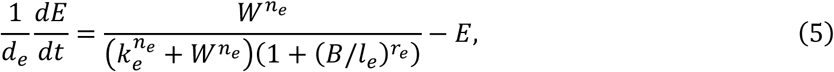

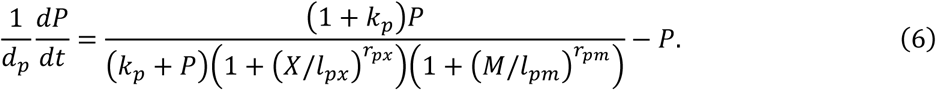

Eq. (1) describes the activation of the BMP4 signalling pathway as a Hill function, which can be interpreted as a generalised Michaelis-Menton kinetics. Eq. (2) describes the diffusion, activation by BMP4 response, and self-activation of WNT/NODAL. We have used the concentration of WNT to represent the overall activation level of the WNT/NODAL pathway, as justified in the main text and the supplementary mathematical information (SM). Eqs. (3-5) govern the evolution of cell fate likelihoods for the three different cell types that we consider: amnion-like (*X*), mesoderm (*M*), and endodermal (*E*). Eq. (6) describes the baseline pluripotent cell state. Eqn. (2) takes into account WNT/NODAL diffusion within the tissue which plays an important role in the gradient chip. Given a known spatio-temporal BMP4 stimulation profile, we simulated the evolution of cell fates in an experimental scenario (gradient chip or µTransWell) by solving the equations numerically using the forward Euler scheme and the standard central finite difference to discretise the spatial derivative of *W*.

The parameters in Eq. (1-6) are provided in table M3 in the SM. The parameters *n*_b_, *k*_b_, *n*_w_, *k*_w_, *n̅*_w_, *k̅*_w_, *a*_w_, *d*_w_, *D*_w_, were obtained from our experimental measurements and data in the previous literature. We fitted the other model parameters **p** = [*n*_x_, *k*_x_, *r*_x_, *l*_x_, *n̅*_x_, *k̅*_x_, *a̅*_x_, *d*_x_, *n*_m_, *k*_m_, *r*_m_, *l*_m_, *d*_m_] against imaging data of cell fate distributions from the microfluidic gradient chip at different BMP4 stimulation strengths. The fitting was based on minimising a cost function measuring the difference between the model predicted and experimentally imaged cell fate profiles at the given time *t*_g_ when measurement was taken:

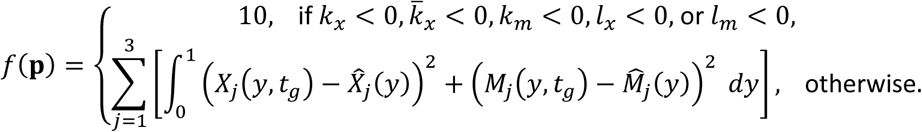

*X̂*_*j*_(*y*), *M̂*_*j*_(*y*) are experimentally identified target cell identity marker distributions, and the subscript *j* (*j* = 1,2,3) denotes three different gradient maximums. To solve the optimisation problem, we adopted the Nelder-Mead method. All numerical calculations were performed with in-house codes written in Julia utilising open-source packages. More details of modelling and extended discussions about our results can be found in the SM. Parameters that are used to produce Fig.5CD are reported in Table M3 in the supplementary. Numerical codes for simulations and analyses are available in the publicly available Github repository (see **Data availability**).

## Data availability

Sample experimental images, Matlab and Python codes to process and analyze the images, and Julia codes for model simulations and analyses are available in the repository: https://github.com/bsorre/wyatt_gradients

## table of primary antibodies used for this study

| Antibody | Organism | Dilution | Supplier | Reference |
| --- | --- | --- | --- | --- |
| SOX2 | Rat | 1:500 | invitrogen | 14-9811-82 |
| OCT3/4 | Mouse | 1:300 | BD Biosciences | 611203 |
| Brachyury/TBXT | Goat | 1:300 | R&D Systems | AF2085 |
| SOX17 | Goat | 1:300 | R&D Systems | AF1924 |
| FOXA2 | Rabbit | 1:250 | Cell Signalling | 8186S |
| pSMAD1/5 | Rabbit | 1:250 | Cell Signalling | 13820 |
| LEF1 | Rabbit | 1:250 | Cell Signalling | 2230S |
| pSMAD2/3 | Rabbit | 1:250 | Cell Signalling | 18338 |
| CDX2 | Mouse | 1:250 | Abcam | ab157524 |
| CDX2 | Rabbit | 1:300 | Abcam | ab76541 |
| ISL1 | Mouse | 1:50 | DSHB | 39.4D5-c |
| GATA3 | Goat | 1:300 | R&D systems | AF2605 |
| TFAP2A | Mouse | 1:50 | santa cruz<br>biotechnologies | sc-12726 |

| **secondary: donkey anti:** |  |  |  |  |
| --- | --- | --- | --- | --- |
| target species | fluorophore | supplier | ref | dilution |
| Mouse | Alexa488 | Abcam | ab150109 | 1:500 |
| Rat | DyLight488 | Thermofischer | SA5-10026 | 1:500 |
| Rabbit | Alexa555 | Abcam | ab150062 | 1:500 |
| Goat | Alexa555 | Abcam | ab150134 | 1:500 |
| Mouse | Alexa647 | Abcam | ab150111 | 1:500 |
| Goat | Alexa647 | Abcam | ab150135 | 1:500 |
| Rabbit | Alexa647 | Abcam | ab150063 | 1:500 |

## Acknowledgements

We thank all members of the teams of Pascal Hersen and Jerome Collignon (IJM, PARIS) for fruitful discussion throughout the project. We thank A.H. Brivanlou for providing the RUES2 and GLR hESC line, the mechanical workshops of the MSC and PCC labs for design and fabrication of custom parts for the gradient chip and Carine Vias at the BMBC platform for help with molecular biology. This work was funded by grants from Agence Nationale de la Recherche (ANR-15-CE13-0007, ANR-23-CE13-0018), CNRS (emergence INC2023), Labex CellInScale, Qlife and Human Frontier Science Program (HFSP CDA00063/2015-C). M. Q. has benefitted from the support of ANR-18-CE13-0028-03 (GuideFusion) attributed to VH

## Author contributions

Conceptualization, TW, MQ, PH, VH, BS; Methodology, TW, SB, PH, BS; Investigation, TW, MQ, JS, HO, SB, BS; Formal Analysis, TW, MQ, JS, GT, MQ, VH, BS; Writing, TW, MQ, JS,VH, BS; Supervision, VH, BS.

