## supplementary figures and legends for "“Morphogen gradients applied basally to human embryonic stem cells to control and dissect tissue patterning”"

### Wyatt et al. Supplementary figures

### Table on content

#### Supplementary figures

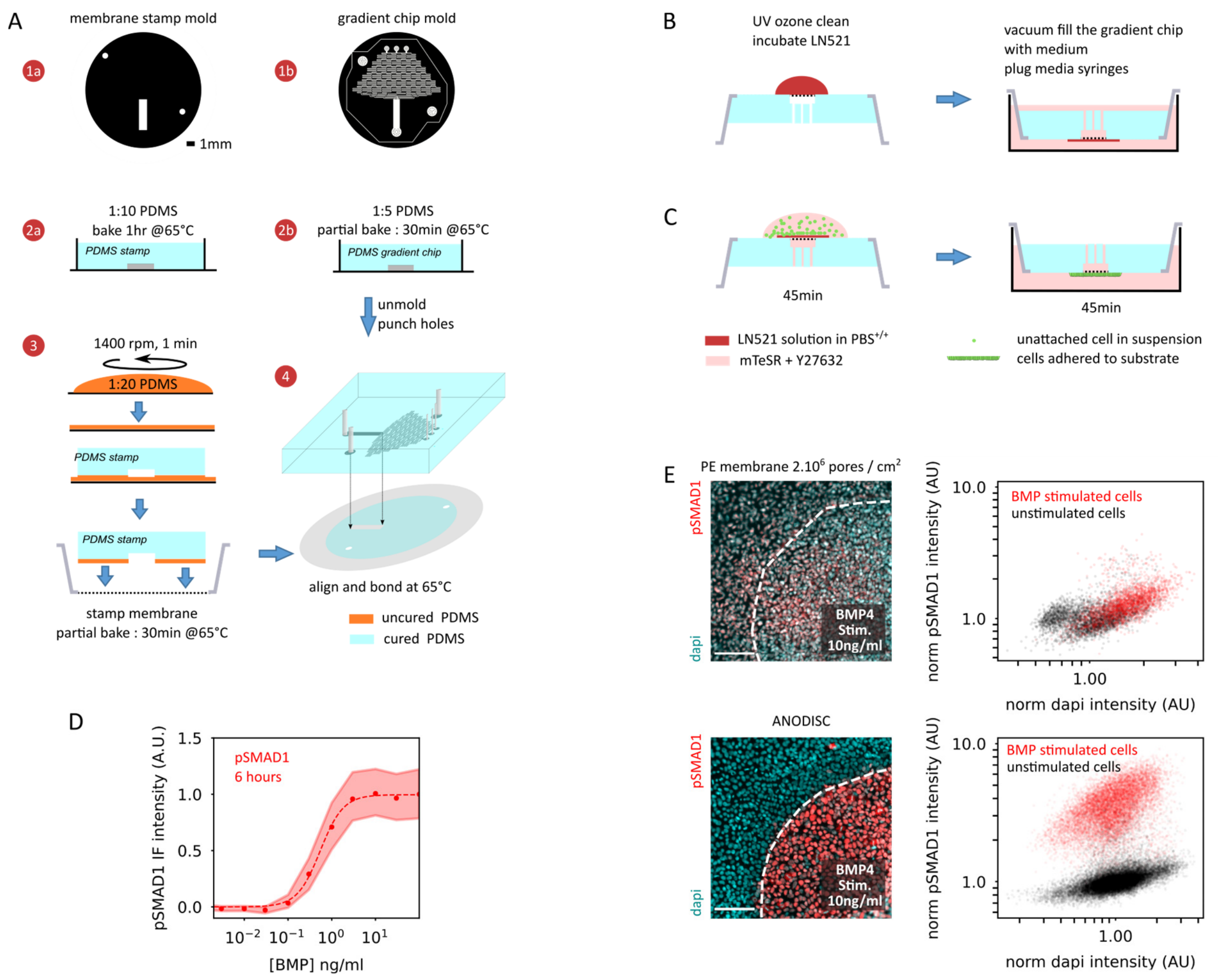

**Figure S1: gradient chip fabrication and characterization of BMP signaling**

**A)** Principle of fabrication of the microfluidic chip to generate morphogen gradient on the basal side of hESC colonies. ***1-*** molds for the membrane stamps (a) and the gradient generator (b) are fabricated on silicon wafers using conventional SU8 negative photoresist. White areas are at 100µm high compared to the reference (black). ***2a-*** PDMS at a 1:10 reticulant:base ratio is poured on the stamp mold and baked for at least 1hour to create the PDMS stamp. After curing, the PDMS block is cut out of the mold. ***2b-*** PDMS at a 1:5 reticulant:base ratio is poured on the gradient chip mold and baked for 30min at 65°C to create a partially cured PDMS gradient chip. After curing, the PDMS block is cut out of the mold and holes for media inlets are punched. ***3-*** 1:20 base reticulant PDMS is spin coated on a glass coverslip for 1 minute at 1400rpm. Then the PDMS stamp from 2a is inked on this thin spin-coated layer of PDMS and then put in contact with an anodisc membrane attached to the frame of a commercial transwell device. PDMS stamped anodisc membrane is then baked for 30 minutes at 65°C. ***4-*** the gradient chip from 2b is aligned with the stamped anodisc membrane so that the gradient chamber faces the rectangle of membrane that is not covered by PDMS. The aligned chip+stamped membrane are then cured for at least one hour to achieve permanent bonding of the 2 elements.

**B)** Principle of chip preparation before cell seeding. First the chip is sterilized 7 minutes on both sides in a UV-Ozone cleaner. Then a drop of Laminin-521 (20µg/ml in PBS$^{+/+}$) is incubated on top of the gradient window for 45 minutes. The chip is then dipped in a well of a 6 well plate full of DMEM/F12 medium and degassed under vacuum for 45 minutes so that medium completely fills the gradient generating circuit. Syringes filled with mTeSR with or without BMP4 are then attached to the chip.

**C)** Principle of cell seeding. A drop of hESC suspension (70µl @ $3.10^6$ cell/ml in mTeSR+Y27632) is incubated for 45 minutes on the upside-down gradient chip to allow cells to attach on top of the gradient window. The chip is then returned and dipped in medium for an extra 45 minutes in mTeSR+Y27632. The chip is then transferred to a custom made imaging cuvette (see main figure 1) filled with mTeSR without Y27632. Stimulation starts usually 6-15 hours after removal of Y27632 by flowing media with a constant flow rate of 12µl/hour with syringe pumps.

**D)** Quantification of the dose-response relationship between pSMAD1 immunofluorescence and BMP4 concentration after 6hours of stimulation for sparse cultures of hESC in glass bottom 96 well plates. Colored areas represent the spread between the 1st and 3rd quartiles of the distribution of the signal of individual cells. Dotted lines represent the fit of the curves with a Hill function:

$pSMAD1 = \frac{[BMP4]^n}{K^n_{pSMAD1}+[BMP4]^n}$. Fitting several replicates yielded the following average parameters for the dose response : $K^{6h}_{pSMAD1} = 0.82 \pm 0.14\, ng/ml,\ n^{6h} = 1.36 \pm 0.1$ , 6 independent replicates, uncertainty is SEM.

**E)** Comparison of the influence of membrane porosity. Left, representative pictures of immunofluorescence pictures of hESC stimulated with 10 ng/ml of BMP4 through 2 types of porous membranes (top: commercial transwell device with Polyester membrane, $10^6$ pores/$cm^2$, corresponding roughly to 1 pore/ cell. Bottom anodisc membrane with about 80% porosity) on each picture pores of the membranes are obstructed on defined areas with PDMS, so that only cells on areas delimited by dotted lines receive BMP4. Right, single cell quantification of the pSMAD1 immunofluorescence for stimulated and unstimulated cells on each membrane, normalized by the average fluorescence of unstimulated cells. Scale Bar : 100µm

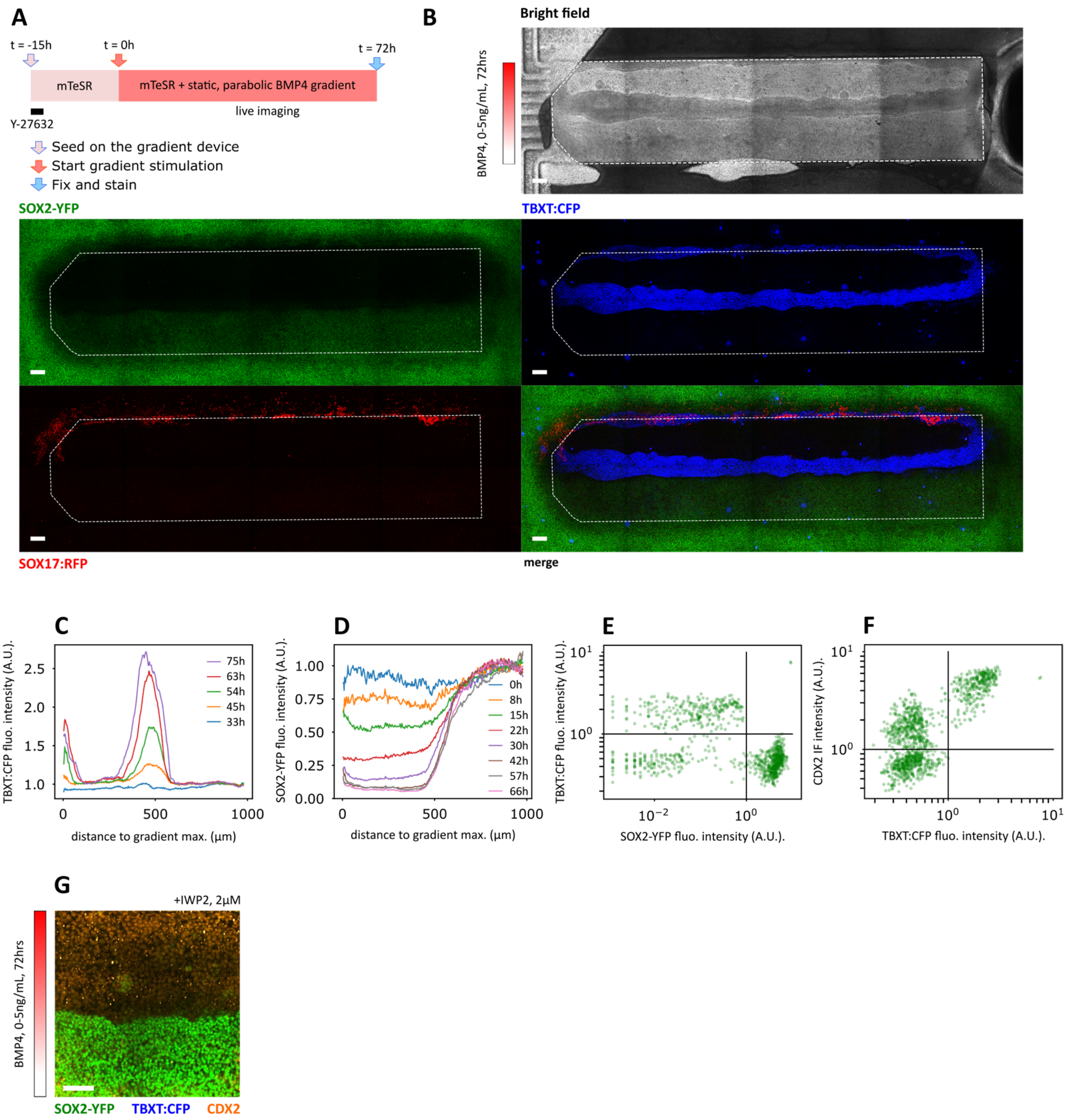

**Figure S2 additional characterizations of differentiation in a static gradient of BMP4**

**A)** experimental timeline of a differentiation experiment under static parabolic gradient of BMP4.

**B)** full scan of the gradient experiment shown in main figure 2. Top: bright field imaging and bottom : fluorescence of the GLR cell line, individual and merged channels.

**C)** temporal evolution of TBXT:CFP during a differentiation experiment under static parabolic gradient with $[BMP4]_{max}$ = 5ng/ml.

**D)** temporal evolution of SOX2-YFP during a differentiation experiment under static parabolic gradient with $[BMP4]_{max}$ = 5ng/ml.

**E,F)** Normalized fluorescence signal of individual cells for the TBXT:CFP vs SOX2-YFP (E) and CDX2 IF vs. TBXT:CFP (F). Horizontal and Vertical full lines represent the thresholds used to define identity for panel D of main figure 2. Undifferentiated cells were defined as [$SOX2^+$/$TBXT^-$/$CDX2^-$], mesoderm cells as [$SOX2^-$/$TBXT^+$/$CDX2^+$],and amnion-like cells as [$SOX2^-$/$TBXT^-$/$CDX2^+$].

**G)** Gradient chip experiment performed following the same timeline as in figure 2 with a static parabolic gradient of $[BMP4]_{max}$ = 10ng/ml but in the presence of the inhibitor of WNT secretion IWP2 (2µM). No $TBXT^{+}$ or $SOX17^{+}$ cells are observed in that condition.

Scale bars: 100µm

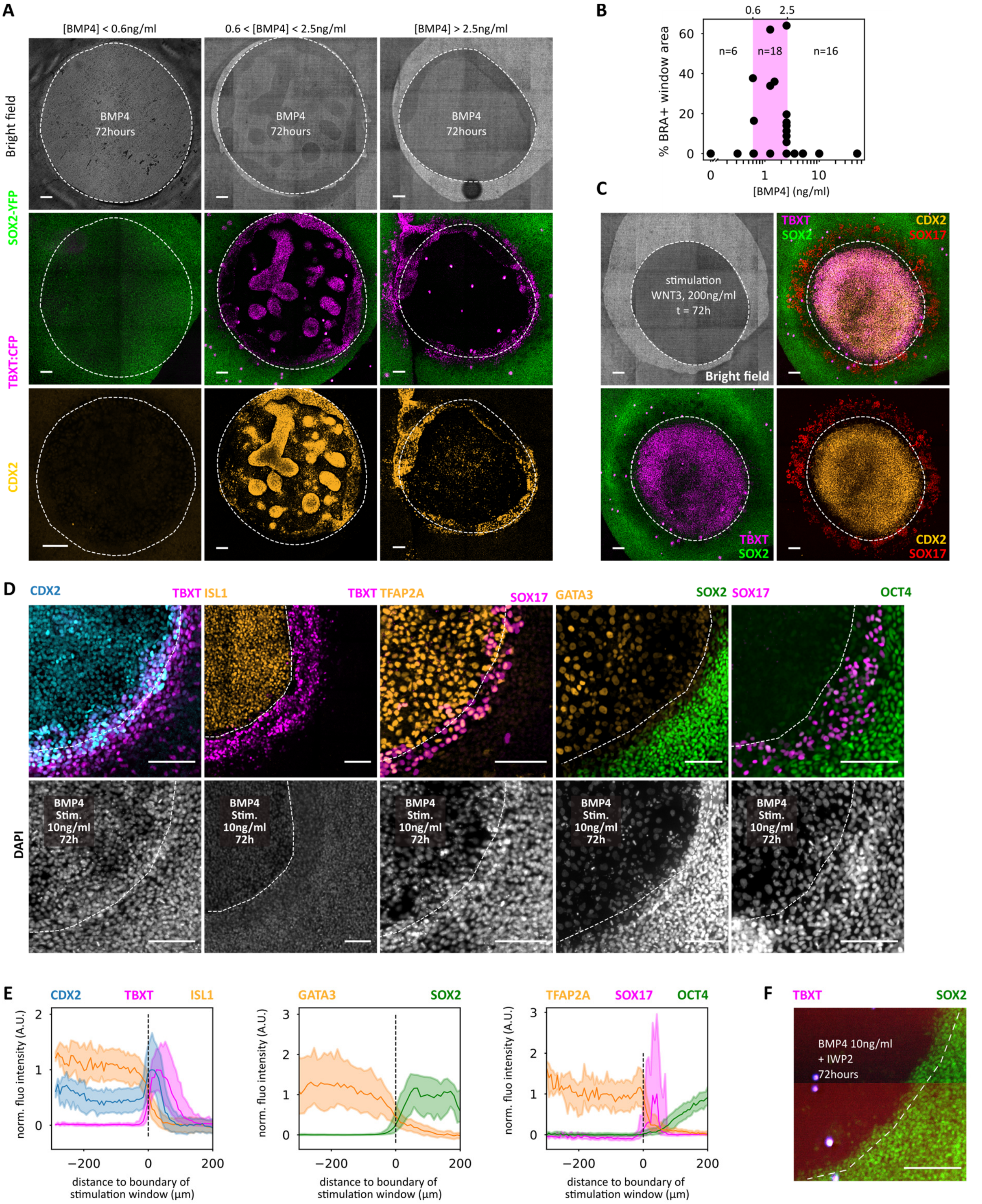

**FigS3 : additional characterization of differentiation on µTransWells.**

**A)** Maximum intensity projections of SOX2-YFP, TBXT:CFP and CDX2 immunofluorescence of a µTransWell experiment after 72 hours of stimulation of the central patch of cells with BMP4 on their basal side at concentration indicated on top of each picture. The dotted line represents the boundary between the stimulated and non-stimulated area of the colony. Experimental timeline is the same as in Fig3. Below 0.6 ng/ml, no differentiation is observed, cells stay [$SOX2^+$/$TBXT^-$]. If [BMP]>2.5 ng/ml, a ring of $TBXT^+$ cells appears at the edge of the stimulation window. Cells on the stimulation window are [$SOX2^-$/$TBXT^-$/$CDX2^+$]. For intermediate doses, (0.6<[BMP4]<2.5ng/ml) patches of $TBXT^+$ cells are observed on the stimulation window, in variable quantity. The

CDX2 stain for the condition [BMP]<0.6ng/ml is from a different experiment than the TBXT:CFP and SOX2-YFP images.

**B)** Quantification of the area covered by patches of $TBXT^+$ cells on the stimulation window as a function of applied BMP4 concentration for n=40 colonies. $TBXT^+$ patches were only observed for 0.6<[BMP4]<2.5ng/ml (n=18/40) and covered at most 60% of the stimulation window.

**C)** Maximum intensity projections of SOX2-YFP, TBXT:CFP, SOX17:RFP and CDX2 immunofluorescence stain of a µTransWell experiment after 72 hours of stimulation of the central patch of cells with 200 ng/ml WNT3 on their basal side. The dotted line represents the boundary between the stimulated and non-stimulated area of the colony. In that case 100% of the cells on the differentiation window are [$SOX2^-$/$TBXT^+$/$CDX2^+$]. Endoderm cells [$SOX17^+$] appear at the edge of WNT-stimulated area. Unlike $SOX17^+$ cells appearing in BMP4-stimulated µWells, these cells are isolated and motile. Representative image of 4 colonies in 2 independent experiments.

**D)** Representative Maximum intensity projections of immunofluorescence of µTransWell experiments after 72 hours of stimulation of the central patch of cells with 10ng/ml BMP4 on their basal side. The dotted line represents the boundary between the stimulated and non-stimulated area of the colony. Cells on the stimulation window are positive for CDX2, ISL1, GATA3 and TFAP2A, a combination consistent with extra-embryonic amnion-like identity. Unstimulated cells away from the stimulation window are positive for pluripotency markers SOX2 and OCT4.

**E)** Average radial profile of expression of immunofluorescence markers displayed in D. Plain lines are the average fluorescence of individual cells from 5 colonies. Colored areas represent the spread between the $1^{st}$ and $3^{rd}$ quartiles of the distribution of the signal of individual cells. Negative distances indicate a position inside the BMP stimulation area.

**F)** µTransWell experiment stimulated for 72hours with 10ng/ml BMP4 supplemented by the inhibitor of WNT secretion IWP2 (2µM). Like in gradient experiments, no $TBXT^+$ or $SOX17^+$ cells are observed in that condition.

Scale bars : 100µm

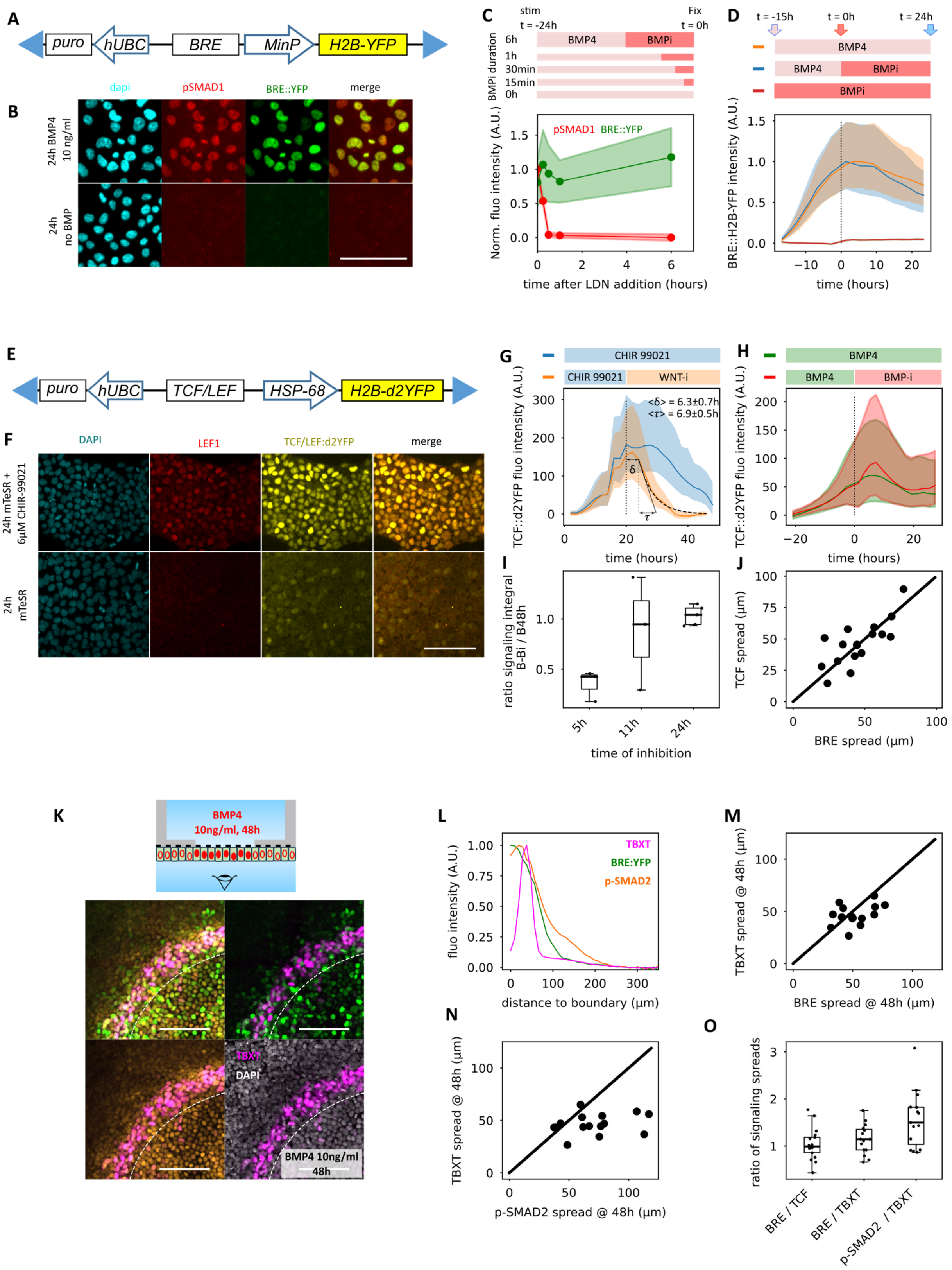

**Figure S4. Characterization of the BRE:YFP and TCF/LEF:d2YFP reporter and additional signaling data after 48h of differentiation in µTransWell experiments.**

**A)** Schematic representation of the transgenic BRE::YFP reporter. The BRE element is upstream a minimal promoter (minP) driving H2B-YFP expression. For image segmentation purposes, the mCherry-NLS nuclear marker is also randomly inserted in the genome with the ePiggyBAC system of transposable elements. Blue triangles represent 5' and 3' terminal repeats required for transposition. Resistance genes are also inserted under the control of hUBC promoter for selection.

**B)** Images showing the induction of phosphorylated SMAD1 (pSMAD1) detected by IF and BRE::YFP after 24hours of stimulation of sparse hESC cells in a well of a 96 well plate with 10ng/ml BMP4. Scale bar = 50µm

**C,D)** comparison of the stability of the pSMAD1 and BRE:YFP reporters. Experimental timelines are displayed on top of each graph. **(C)** 30minutes after addition of the BMP pathway inhibitor LDN ($BMP_i$) pSMAD1 is not detected by immunofluorescence while BRE:YFP fluorescence level remains unchanged even after 6hours. (**D**) long term inhibition experiment shows that after 24 hours of BMP inhibition, BRE:YFP signal is only reduced by 15% compared to control. Shaded areas represent the $1^{st}$ and $3^{rd}$ quartiles of the distribution of individual cells.

**E)** Schematic representation of the transgenic TCF/LEF:d2YFP reporter. The TCF/LEF element is upstream a minimal promoter (hsp68) driving expression of a destabilized YFP linked to a histone for nuclear localization (H2B-d2YFP). Blue triangles represent 5' and 3' terminal repeats required for transposition. A resistance gene (puro) is also inserted under the control of hUBC promoter for selection.

**F)** Images showing the induction of LEF1, a transcriptional target of the WNT/βcat pathway detected by IF and of the TCF/LEF::d2YFP reporter after 24hours of stimulation of sparse hESC cells in a well of a 96 well plate with 6µM of the WNT agonist CHIR99021. Scale bar = 50µm

**G)** Measurement of the stability of the TCF/LEF:d2YFP reporter according to the experimental timelines displayed on top of the graph. After 20 hours of stimulation, the medium containing the WNT/βcat pathway agonist CHIR99021, is replaced by medium containing inhibitors IWP2 and XAV. Unlike the BRE:YFP reporter, the fluorescence signal of the TCF/LEF:d2YFP reporter decays faster in inhibited cells compared to the control. After a lag time $\delta = 6.3 \pm 0.7h$ , the decay of the fluorescence signal is well fitted by a single exponential decay $TCF = Ae^{-t/\tau}$ with $\tau = 6.9 \pm 0.5\ h$, n=6 independent replicates, uncertainty is s.e.m.

**H)** BMP stimulation induces an increase in WNT/β-cat pathway activity as revealed by an increase of fluorescence of TCF/LEF:d2YFP reporter (green curve). However, blocking BMP signaling with the BMP inhibitor LDN after 20 hours of stimulation (red curve) does not trigger a decrease of the TCF/LEF:d2YFP reporter fluorescence, suggesting that WNT signaling can sustain itself without the need of an active BMP pathway.

**I)** Estimation of the minimum duration of BMP stimulation necessary to induce self-sustained WNT/β-cat signaling by quantification of the ratio of integral of TCF/LEF:d2YFP signaling (area under the curve) for constant BMP4 stimulation for 48h versus transient BMP4 stimulations during 5, 11 or 24 hours, see timeline of panel D. 5 hours of BMP stimulation are not sufficient to trigger self-sustained WNT/β-cat signaling. 24 hours of stimulation are always sufficient to induce TCF/LEF:d2YFP signaling at least as high as the control. Outcome is more variable for 11 hours stimulation, suggesting that this time is close to the threshold duration necessary to induce self-sustained WNT/β-cat signaling. Each dot represents the ratio value of an independent replicate.

**J)** Spread of BRE:YFP vs. TCF/LEF:d2YFP signal after 24h of BMP stimulation in µtransWell experiments reported in main figure 4H. Each dot represents the measurement of an independent replicate. The full line (y=x) is added as a guide to the eye and is not a fit of the data.

**K)** Top: schematic reminder of the µTransWell configuration. Bottom: BRE:YFP, pSMAD2 and TBXT immunofluorescence of a µTransWell experiment after 48 hours of stimulation of the central patch of cells with BMP4 (10ng/ml) on their basal side. The dotted line represents the boundary between the stimulated and non-stimulated area of the colony. Scale bar = 100µm

**L)** Radial profiles of expression of BRE:YFP, pSMAD2 and TBXT as a function of the distance to the boundary of the stimulation window of a µTransWell experiment after 48 hours of stimulation of the central patch of cells with BMP4 (10ng/ml). Curves are average of single cell signal for 4 independent replicates

**M)** Spread of TBXT vs. BRE:YFP signal after 48h of BMP stimulation in µTransWell experiments. Each dot represents the measurement of an independent replicate. The full line (y=x) is added as a guide to the eye and is not a fit of the data.

**N)** Spread of TBXT vs. pSMAD2 signal after 48h of BMP stimulation in µTransWell experiments. Each dot represents the measurement of an independent replicate. The full line (y=x) is added as a guide to the eye and is not a fit of the data.

**O)** Box plots representing the ratio of measured spreads of signaling outside the stimulation area (as defined in Fig4B) for indicated pairs of signaling and fate reporters. Dots represent individual replicates. Signaling reporters (TCF/LEF:d2YFP, BRE:YFP, pSMAD2,) and TBXT spread have been assayed after 48 hours of BMP stimulation.

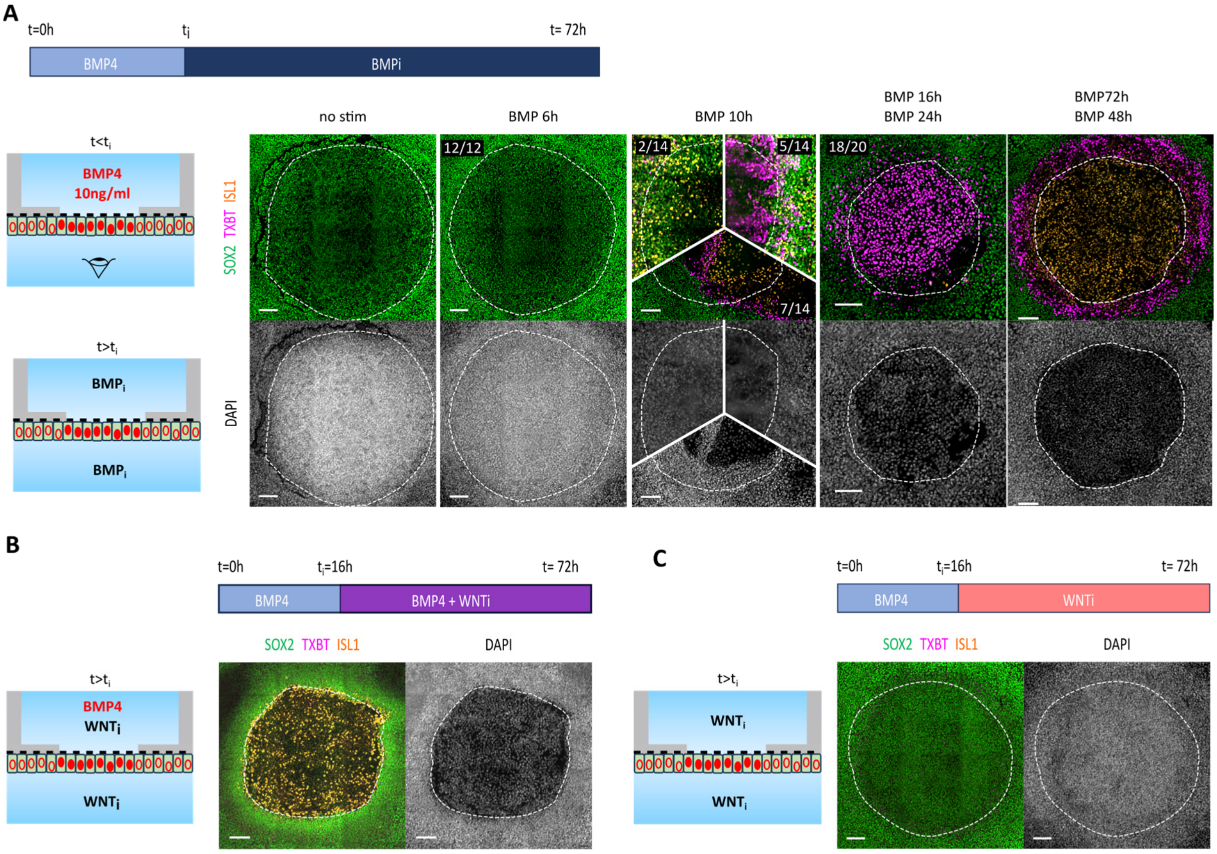

**Figure S5. Effect of additional regimen of signaling inhibition on differentiation and patterning.**

**A)** Maximum intensity projections of SOX2, TBXT and ISL1 immunofluorescence after 72 hours of differentiation in a µTransWell device for different durations of BMP4 stimulation. Initially, a central patch of cells is stimulated with BMP4 (10ng/ml) on their basal side. The dotted line represents the boundary between the stimulated and non-stimulated area of the colony. At time $t_i$, indicated on top of individual pictures, the basal BMP stimulation is replaced by inhibition with the BMP inhibitor LDN. When BMP is inhibited after 16 or 24 hours ISL1$^+$ amnion-like cells observed on the stimulation window in the control (BMP 72h) are replaced by TBXT$^+$ cells (18/20 colonies over 5 independent experiments). 6 hours of BMP stimulation are not sufficient to induce differentiation (12/12 colonies over 3 independent experiments). For intermediate duration of BMP stimulation (10h) the outcome was variable. Over 14 colonies in 3 independent experiments, 2 colonies were composed of SOX2+ cells with sparse ISL1+ cells and no TBXT$^+$ cells. The remaining colonies had either differentiated cells on the stimulation window, a majority of TBXT$^+$ cells (5/14 colonies) or patches of TBXT$^+$ and ISL1$^+$ cells (7/14).

**B,C)** Maximum intensity projections of SOX2, TBXT and ISL1 immunofluorescence after 72 hours of differentiation in a µWell device. During the first 16 hours of experiments, a central patch of cells is stimulated with BMP4 (10ng/ml) on their basal side. The dotted line represents the boundary between the stimulated and non-stimulated area of the colony. 16 hours after the initiation of BMP stimulation, inhibitors of WNT/β-cat signaling activity (XAV+IWP2) were added to the apical compartment of the device and the basal medium was replaced by BMP4 + WNT inhibitors (B) or WNT inhibitors alone (C). In both cases, the ring of TBXT$^+$ cells at the edge of the BMP-stimulated area observed in the control (see panel A) disappeared. ISL1 was expressed by cells on the stimulation window only in colonies where BMP stimulation was sustained. Representative pictures of 10 colonies over 2 independent experiments.

Scale bars are 100µm.

**Description of supplementary movies**

**Supplementary movie 1**: time lapse movie of filling of the gradient chip with media containing fluorescent dextrans.

**Supplementary movie 2**: time lapse of differentiation of GLR hESC under stimulation with a static parabolic gradient of BMP4 with $[BMP4]_{max}$ = 5 ng/ml

**Supplementary movie 3:** Z stack showing the 3D organization of the SOX17+ and TBXT cells at the edge of the stimulation window on a BMP4 stimulated µTransWell experiment

**Supplementary movie 4:** time lapse movie showing the differentiation of GLR hESC on a one BMP4 and one WNT3A stimultated µTranswell experiments

**Supplementary movie 5:** single cell tracking of $BRE:YFP^{+}$ cells found outside of the stimulation window of a µTransWell after 24 hours of BMP4 stimulation
