## Supplementary material for "“Morphogen gradients applied basally to human embryonic stem cells to control and dissect tissue patterning”": mathematical supplement

#### 1 Introduction

We have developed a model for the cell fate network governing the patterning of human Embryonic Stem Cells (hESCs) in our experiments, coupled to the signalling of WNT driven by the morphogen BMP4. Similar to Camacho-Aguilar *et al.*[1], our model aims to capture in a simplified way the interactions between cell fates rather than attempting to describe the full regulatory network underlying these fate choices. The latter is usually complex involving a large number of only partially known biochemical species, which would both make its description difficult and increase the number of unknown parameters.

We consider the cell fate network of Fig. 5B. Stimulation by the morphogen BMP4 induces directly the extra-embryonic, amnion-like fate (CDX2$^+$/GATA3$^+$/ISL$^+$/TFAP2A$^+$ and TBXT$^-$)[2–5]. The mesoderm fate (CDX2$^+$/TBXT$^+$) is induced indirectly via BMP4-initiated production of WNT and NODAL[6–10](Fig. S2, S3). These two secreted morphogens can both diffuse and self-activate[11–13]. As a simplification, here WNT and NODAL are considered to be a single entity. In $\mu$TransWells experiments, expression of TBXT coincides with the domain where WNT signalling is active while NODAL signalling is activated further away from the BMP4 stimulation window without inducing TBXT expression (Figs. 4&S4). Activation of WNT signalling is thus a limiting factor for TBXT expression and therefore, we designate the WNT/NODAL entity as WNT in the current model. The same simplification has also been used recently in Camacho-Aguilar *et al.*[1] Considering WNT and NODAL as separate entities would be necessary, however, to model the binary choice between mesoderm and endoderm fates[14], which is out of the scope of the present study. The amnion-like and mesoderm fates cross repress each other, consistent with their mutually exclusive cell identities.

We further consider a baseline pluripotent state (SOX2$^+$/OCT4$^+$/CDX2$^-$/TBXT$^-$) (Fig. S3) which describes the initial state of the embryonic stem cells. It is self-maintaining and cross inhibits with the amnion-like and mesoderm fates. In addition, we also account for the endoderm fate (SOX17$^+$/FOXA2$^+$)[15] which is induced by WNT and NODAL signalling but prevented by BMP4[8,16]. In this simplified description, since the endodermal cells are downstream of BMP4/WNT dynamics and are only found outside the BMP4 stimulation windows, we neglect the cross inhibition of the endoderm fate with other cell states. The cell types that are considered in this study together with their combinatorial expression of main identity markers are summarised in Table M1.

| cell type | CDX2 | ISL1 | TBXT | SOX17 | FOXA2 | SOX2 | OCT4 |
|---|---|---|---|---|---|---|---|
| extra-embryonic/amnion-like (X) | + | + | - | - | - | - | - |
| mesodermal (M) | + | - | + | - | - | - | - |
| endodermal (E) | - | - | - | + | + | - | - |
| pluripotent (P) | - | - | - | - | - | + | + |

Table M1: Cell fates characterised by the expression of identity markers.

### 2 Model

#### 2.1 Formulation

Let $\tilde{b}$ be the externally supplied BMP4 concentration. We define the BMP4 pathway activity level using another variable $\tilde{B}$ as a measure of the cells' response, which could be interpreted as the nuclear concentration of pSmad1/5 that can be measured experimentally (Fig. 1I and S1). $\tilde{W}$ denotes the activity level of the WNT/$\beta$-catenin pathway. The variables $\tilde{X}, \tilde{M}, \tilde{E}$ represent the *likelihoods* of cells differentiating into extra-embryonic, mesodermal, and endodermal tissues, respectively, while $\tilde{P}$ denotes the likelihood for staying pluripotent. $\tilde{X}, \tilde{M}, \tilde{E}$ and $\tilde{P}$ can be interpreted as proxies for probabilities of adopting the corresponding cell types. We do not, however, impose any normalisation condition since these are not actual probabilities and we neglect other cell identities. Given the cell fate network in Fig. 5B, we consider the following governing equations. First, pSmad1/5 is activated by BMP4 and represents the system's response to stimulation

$$\tilde{B} = \frac{A_b \tilde{b}^{n_b}}{K_b^{n_b} + \tilde{b}^{n_b}}, \tag{M1}$$

where $n_b$ is the Hill coefficient, $K_b$ the half-saturation constant, and $A_b$ the activation strength. Second, WNT is diffusive and induced by cells in response to BMP4. In addition, we assume that WNT is self-activating, which is critical for sustained WNT activity after transient BMP4 stimulation in the $\mu$TransWells (Figs. 4, 5E-H, S4 and S5, further detailed in Sec. 3.1). We only consider one spatial dimension because cell fate variation perpendicular to the BMP4 gradient is much smaller than along the gradient (Figs. 2 and S2).

$$\frac{\partial \tilde{W}}{\partial T} = \eta_w \frac{\partial^2 \tilde{W}}{\partial Y^2} + \frac{A_w \tilde{B}^{n_w}}{K_w^{n_w} + \tilde{B}^{n_w}} + \frac{\bar{A}_w \tilde{W}^{\bar{n}_w}}{\bar{K}_w^{\bar{n}_w} + \tilde{W}^{\bar{n}_w}} - \delta_w \tilde{W}, \tag{M2}$$

where $\eta_w$ is an effective intercellular diffusivity. $(n_w, K_w, A_w)$, $(\bar{n}_w, \bar{K}_w, \bar{A}_w)$ are constants associated with the Hill functions describing WNT activation by BMP4 and itself, respectively. $T$ denotes time, and $Y$, as the spatial coordinate, is the distance along the gradient from the maximum BMP4 concentration. The last term represents the decay of WNT with a constant rate $\delta_w$. The evolution of the fate markers for the amninon-like ectoderm, mesoderm and endoderm is governed by

$$\frac{d\tilde{X}}{dT} = \left( \frac{A_x \tilde{B}^{n_x}}{K_x^{n_x} + \tilde{B}^{n_x}} + \frac{\bar{A}_x \tilde{X}^{\bar{n}_x}}{\bar{K}_x^{\bar{n}_x} + \tilde{X}^{\bar{n}_x}} \right) \frac{1}{\left(1 + (\tilde{M}/L_x)^{r_x}\right)\left(1 + (\tilde{P}/L_{xp})^{r_{xp}}\right)} - \delta_x \tilde{X}, \tag{M3}$$

$$\frac{d\tilde{M}}{dT} = \frac{A_m \tilde{W}^{n_m}}{\left(K_m^{n_m} + \tilde{W}^{n_m}\right)\left(1 + (\tilde{X}/L_m)^{r_m}\right)\left(1 + (\tilde{P}/L_{mp})^{r_{mp}}\right)} - \delta_m \tilde{M}, \tag{M4}$$

$$\frac{d\tilde{E}}{dT} = \frac{A_e \tilde{W}^{n_e}}{K_e^{n_e} + \tilde{W}^{n_e}} \frac{1}{1 + (\tilde{B}/L_e)^{r_e}} - \delta_e \tilde{E}. \tag{M5}$$

In Eqn. M3, the two Hill functions on the right hand side (r.h.s.) represent activation of the amnion-like fate by BMP4 and its self-activation, respectively, while the inhibitory Hill functions describe repression of the this fate by the mesodermal and pluripotent cells, respectively. $(n_x, K_x, A_x)$, $(\bar{n}_x, \bar{k}_x, \bar{A}_x)$, $(r_x, L_x)$, and $(r_{xp}, L_{xp})$ are constants associated with these Hill functions. Here we have accounted for the inhibition of the extra-embryonic, amnion-like identity $X$ by the pluripotent $P$ and mesodermal $M$ tissues through multiplying consecutive inhibitory Hill functions because as $P, M \to 0$, inhibitory effects should vanish. In Camacho-Aguilar *et al.*[1], the authors have adopted a slightly different form assuming that the inhibitory effects are additive rather than multiplicative, but we expect similar qualitative behaviour in both models. $\delta_x$ is a constant decay rate of the amnion-like fate marker $\tilde{X}$. The mesoderm fate $\tilde{M}$ is activated by WNT with Hill parameters $(n_m, K_m, A_m)$, inhibited by the extra-embryonic fate $\tilde{X}$ with Hill parameters $(r_m, L_m)$ and by the baseline pluripotent state $\tilde{P}$ with Hill parameters $(r_{mp}, L_{mp})$, and decaying at the constant rate $\delta_m$ (Eqn. M4). The endoderm fate $\tilde{E}$ is activated by WNT with Hill constants $(n_e, K_e, A_e)$, inhibited by BMP4 $\tilde{B}$ with Hill parameters

$(r_e, L_e)$, and decaying at rate $\delta_e$ (Eqn. M5). Again, since experimental data only require the endoderm to be co-existent with the mesoderm in the absence of BMP4, we neglect its cross repression with other fates.

Since the pluripotent identity is a baseline state, $\tilde{P}$ dynamics is supposed to have a single stable fixed point in the absence of inhibition by other fates. Therefore, we describe its self activation with a Hill coefficient of unity, together with suppression by the amnion-like fate $\tilde{X}$ (inhibitory Hill parameters $(r_{px}, L_{px})$) and by the mesoderm fate $\tilde{M}$ (inhibitory Hill parameters $(r_{pm}, L_{pm})$), as well as exponential decay at rate $\delta_p$. Eqn. M6 governs the evolution of $\tilde{P}$

$$\frac{d\tilde{P}}{dT} = \frac{(A_p + \delta_p K_p)\,\tilde{P}}{\left(K_p + \tilde{P}\right)\left(1 + (\tilde{X}/L_{px})^{r_{px}}\right)\left(1 + (\tilde{M}/L_{pm})^{r_{pm}}\right)} - \delta_p \tilde{P}. \tag{M6}$$

$K_p$ and $A_p$ are constants related to the activating Hill function. Clearly when $\tilde{X} = \tilde{M} = 0$, $\tilde{P}$ tends to the reference value $A_p/\delta_p$ in time, as desired.

To nondimensionalise the system, we introduce characteristic length and time scales $L_0,\ t_0$, respectively, which will be specified later. Suppose there is a characteristic BMP4 concentration $b_0$ whose value is not necessary to be known. Define dimensionless concentrations and likelihoods

$$b = \frac{\tilde{b}}{b_0},\ B = \frac{\tilde{B}}{A_b},\ W = \frac{\delta_w \tilde{W}}{\bar{A}_w},\ X = \frac{\delta_x \tilde{X}}{A_x},\ M = \frac{\delta_m \tilde{M}}{A_m},\ E = \frac{\delta_e \tilde{E}}{A_e},\ P = \frac{\delta_p \tilde{P}}{A_p}.$$

Here we have scaled the WNT activity level by its characteristic value determined by self-activation as WNT bistability is essential (Figs. 4, 5 and the main text). The extra-embryonic amnion-like likelihood is scaled by its activation through BMP4, while the mesoderm and endoderm fate likelihoods are scaled based on induction by WNT. The pluripotent likelihood is normalised by its own activation since it is the baseline fate. Using dimensionless space $y = Y/L_0$ and time $t = T/t_0$, the dimensionless system is

$$B = \frac{b^{n_b}}{k_b^{n_b} + b^{n_b}}, \tag{M7}$$

$$\frac{1}{d_w}\frac{\partial W}{\partial t} = D_w \frac{\partial^2 W}{\partial y^2} + \frac{a_w B^{n_w}}{k_w^{n_w} + B^{n_w}} + \frac{W^{\bar{n}_w}}{\bar{k}_w^{\bar{n}_w} + W^{\bar{n}_w}} - W, \tag{M8}$$

$$\frac{1}{d_x}\frac{dX}{dt} = \left(\frac{B^{n_x}}{k_x^{n_x} + B^{n_x}} + \frac{\bar{a}_x X^{\bar{n}_x}}{\bar{k}_x^{\bar{n}_x} + X^{\bar{n}_x}}\right)\frac{1}{(1 + (M/l_x)^{r_x})\,(1 + (P/l_{xp})^{r_{xp}})} - X, \tag{M9}$$

$$\frac{1}{d_m}\frac{dM}{dt} = \frac{W^{n_m}}{(k_m^{n_m} + W^{n_m})\,(1 + (X/l_m)^{r_m})\,(1 + (P/l_{mp})^{r_{mp}})} - M, \tag{M10}$$

$$\frac{1}{d_e}\frac{dE}{dt} = \frac{W^{n_e}}{(k_e^{n_e} + W^{n_e})\,(1 + (B/l_e)^{r_e})} - E, \tag{M11}$$

$$\frac{1}{d_p}\frac{dP}{dt} = \frac{(1 + k_p)\,P}{(k_p + P)\,(1 + (X/l_{px})^{r_{px}})\,(1 + (M/l_{pm})^{r_{pm}})} - P, \tag{M12}$$

with the dimensionless groups:

$$k_b = \frac{K_b}{b_0},\ d_w = \delta_w t_0,\ D_w = \frac{\eta_w}{\delta_w L_0^2},\ k_w = \frac{K_w}{A_b},\ a_w = \frac{A_w}{\bar{A}_w},\ \bar{k}_w = \frac{\delta_w \bar{K}_w}{\bar{A}_w},\ \bar{a}_x = \frac{\bar{A}_x}{A_x},\ k_x = \frac{K_x}{A_b},$$

$$\bar{k}_x = \frac{\delta_x \bar{K}_x}{A_x},\ l_x = \frac{\delta_m L_x}{A_m},\ l_{xp} = \frac{\delta_p L_{xp}}{A_p},\ d_x = \delta_x t_0,\ k_m = \frac{\delta_w K_m}{\bar{A}_w},\ l_m = \frac{\delta_x L_m}{A_x},\ l_{mp} = \frac{\delta_p L_{mp}}{A_p},$$

$$d_m = \delta_m t_0,\ k_e = \frac{\delta_w K_e}{\bar{A}_w},\ l_e = \frac{L_e}{A_b},\ d_e = \delta_e t_0,\ k_p = \frac{\delta_p K_p}{A_p},\ l_{px} = \frac{\delta_x L_{px}}{A_x},\ l_{pm} = \frac{\delta_m L_{pm}}{A_m},\ d_p = \delta_p t_0.$$

We have summarised the definitions and biological meaning of these nondimensional parameters in Table M3.

We use the model to study cell differentiation in the two different experimental devices: the BMP gradient chip and the $\mu$TransWell. Given the setups, it is convenient to choose the size of the gradient chip $L_0 = 1\ mm$ as the length scale, which is also on the same order of magnitude as the diameter ($0.7\ mm$) of the stimulation window in the $\mu$TransWell device. We pick the time scale to be $t_0 = 11$ hours, which is the threshold BMP4 stimulation time to induce WNT bistability as measured in the $\mu$TransWell experiments (Figs. S4, S5), as well as the average time spent on the stimulation window by the BRE:YFP$^+$ cells found outside the window in the $\mu$TransWells (Fig. 4D). A more detailed discussion about this time scale is in Sec. 3.1. For the gradient chip, we simulate the transient cell differentiation along a pre-established static BMP4 gradient. In this case we prescribe the BMP4 concentration to be $b(y) = q(1 - y)^2$ on $y \in [0, 1]$, where $q$ is the maximum BMP4 concentration in the gradient and $y$ is the distance to the maximum concentration. For the $\mu$TransWell, we neglect spatial dependence and apply a constant BMP4 concentration $b = b_s$ during a prescribed time period $t_s$ after which BMP4 is removed. This models the process of an individual cell migrating from inside the BMP4 stimulation window to outside as in our experiments. It can also be interpreted as subjecting the cell to sequential uniform BMP4 stimulation and inhibition. Mathematically, compared to the gradient chip simulation, the only partial differential equation Eqn. (M8) reduces to an ordinary differential equation in time, and thus all governing equations are local. The model is parametrised using a combination of these two types of simulations, informed by data from our experiments and literature, as detailed in Sec. 3.

#### 2.2 Numerical solution

In either types of simulations, we must solve the governing equations as initial value problems or initial boundary value problems. Notice that the BMP4 response level $B$ can be readily calculated with the prescribed BMP4 concentration $b$ through Eqn. (M7), while the likelihood of the endoderm fate $E$ is slaved to $B$ and $W$ through Eqn. (M11). Hence these two equations are effectively decoupled from the rest. We solve the coupled Eqns. (M8)–(M10) and (M12) for $W,\ X,\ M,\ P$ together by marching in time from an initial condition of $W = 0,\ X = 0,\ M = 0,\ P = 1$. To simulate the gradient chip which is subject to WNT diffusion across the stimulation window boundaries, we pad each end of the stimulation window $[0, 1]$ with a region of length $l_w = 6\sqrt{D_w}$, where $\sqrt{D_w}$ is the dimensionless diffusion length. Then at both ends of the computational domain, we impose the boundary condition $W = 0$ to mimic vanished WNT concentration at infinity for Eqn. (M8). The spatial derivative is discretised with the standard second-order central finite difference scheme on a computational grid of mesh size $h \approx 1/128$. This grid size is only approximate because it is slightly different inside and outside the window to ensure that one grid point collocates with each of the stimulation window boundaries. For time marching, we use the forward-Euler scheme for its fast speed and low memory requirement thanks to the one-dimensional model. To ensure numerical stability, the time step size is chosen to be $k = k_0/\max\{d_w, d_x, d_m, d_p, 1\}$ where $k_0 = 5 \times 10^{-6}/D_w$ if $D_w \neq 0$, or $k_0 = 2.5 \times 10^{-3}$ if $D_w = 0$. Since the fate likelihoods $X,\ M,\ P,\ E$ are always positive, during time stepping we have implemented a projection scheme that truncates any negative $X,\ M,\ P,\ E$ values and resets them to $10^{-8}$. When modelling the $\mu$TransWell, spatial dependence is neglected, and we simply march Eqns. (M8)–(M10) and (M12) in time. Once the solution is obtained, using Eqn. (M11) we solve for the evolution of the endoderm fate through forward-Euler with the same time steps again. The numerical simulations are implemented using in-house codes in `Julia 1.11.6`[17].

### 3 Parameter fitting

#### 3.1 Preliminaries

At this stage, our aim is to use key experimental insight and measurements to constrain and preset a number of parameters. Firstly, the response of the tissue to BMP4 stimulation through pSmad1/5 has been characterised by experimental data (Figs. 1I, S1G), and $n_b = 1.355$ and $k_b = 0.8233$. Secondly, in our model the BMP4 response and WNT dynamics are upstream to all cell fate choices, which allows us to parametrise WNT dynamics independent of downstream species. We fix $n_w = 4$ and $\bar{n}_w = 6$ so as to have generally sharp profiles of the Hill functions for WNT activation by BMP4 and its self-activation. Small alteration of these values is not expected to result in major changes in the results.

To further determine $\bar{k}_w$ for WNT self-activation, we note that there have been reports of the presence of $\beta$-catenin at the Wnt3 promoter at the onset of primitive streak formation at E6.5 mouse embryo[11], and of WNT self activation in BMP stimulated hESCs[1,13]. In addition, our own experiments provide two lines of evidence supporting that WNT has a bistable behaviour. On one hand, the signalling activity of the WNT/$\beta$-cat pathway is sustained after BMP4 inhibition (Fig. S4), as well as after cells migrating outside the stimulation window when BMP4 induction has stopped (Fig. 4). On the other hand, the inhibition experiment reported in Fig. 5E-H shows that sustained WNT activity after the cells have left the stimulation area is necessary for TBXT expression and thus mesoderm specification. Clearly, after BMP4 stimulation has terminated, the self-activation of WNT must allow for bistability. In this case, for a $\mu$TransWell at steady state, Eqn. (M8) reduces to

$$\frac{W^{\bar{n}_w}}{\bar{k}_w^{\bar{n}_w} + W^{\bar{n}_w}} - W = 0. \tag{M13}$$

Given $\bar{n}_w = 6$, the requirement of bistability imposes an upper bound for the parameter $\bar{k}_w$ at 0.6373 (dashed curve in Fig. M1A), and we pick $\bar{k}_w = 0.56$ (red solid curve in Fig. M1A). The choice of $\bar{n}_w$ and $\bar{k}_w$ could affect the location where WNT transitions from low to high values and thus the fate boundary between the pluripotent and mesoderm cells, to be detailed in Sec. 4.4.

Our data from inhibition experiments, either with WNT activity reporters (Fig. S4I) or with fate outcomes (Fig. S5A), also show that 6 hours of BMP4 stimulation was not enough to induce a self-sustained WNT response or the mesodermal fate, while 16 hours was sufficient. Stimulation around 10-12 hours yielded variable results, suggesting that the threshold time lies in this range. This is consistent with the average time spent by cells on the stimulation window before being pushed out in the $\mu$TransWell experiments (Fig. 4D). It implies that a minimal duration of BMP4 stimulation time of about 11 hours is required for WNT to cross the bistability threshold and become self-sustaining. In our model, this time scale is controlled by $d_w$ ($\delta_w$ in the dimensional formulation) and we thus set $d_w = 1$ ($1/\delta_w = t_0 = 11$ hours). This is consistent with the WNT degradation time scale of approximately 14 hours as inferred in the recent work by Camacho-Aguilar et al.[1], but is much longer than the time scale of 50 minutes considered in an earlier study[13]. A discussion on the impact of this parameter is provided in Sec 4.2.

The threshold BMP4 stimulation time of 11 hours ($t_c = 1$ in nondimensional quantities) further allows us to determine parameters of WNT induction by BMP4. For the $\mu$Transwells, we set $b_s = 10$ as in the experiments. We first notice that in presence of BMP, at steady state Eqn. (M8) becomes

$$\frac{a_w B^{n_w}}{k_w^{n_w} + B^{n_w}} + \frac{W^{\bar{n}_w}}{\bar{k}_w^{\bar{n}_w} + W^{\bar{n}_w}} - W = 0. \tag{M14}$$

Eqn. (M14) must yield a unique solution of $W$ that also needs to be in the basin of attraction of the high WNT level enabled by its bistability. This criterion gives an upper bound of allowable $k_w$ value for each $a_w$ (red curve in Fig. M1B). For particular $k_w,\ a_w$ values, we simulate by solving Eqn. (M8) the long-term WNT state in the $\mu$TransWell as a function of different durations of BMP4 exposure, and discern the critical duration $t_c$ when WNT activity increases past the threshold to evolve towards the high steady-state level, i.e., enabling a stable mesoderm fate after BMP4 stops. We identify the $k_w,\ a_w$ combinations that give $0.75 < t_c < 1.25$ (yellow round markers in Fig. M1B). As expected, these parameters are within the aforementioned validity region for $k_w$. From a range of compatible parameter values, we pick $k_w = 0.96,\ a_w = 1.5$ (star marker in Fig. M1B). Given any combination of $\bar{n}_w,\ \bar{k}_w$ values, we must also check that the chosen $k_w, a_w$ pair ensures that the high WNT level is stable at the target pluripotent/mesoderm fate boundary in the BMP gradient. This will be elaborated in Sec. 4.4 and here the picked parameters satisfy this condition (green solid curve in Fig. M1A).

We then consider WNT diffusion in the BMP4 spatial gradients. We are not aware of any measurement of extracellular WNT diffusivity in early human embryogenesis. For *Xenopus* embryos, Mii et al. reported an apparent WNT diffusivity close to $0.04\ \mu m^2/s$[18]. In our experimental data, at the high concentration end of the BMP4 gradient, the size of the mesoderm- and endoderm-differentiated region outside the stimulation window is below $50\ \mu m$ (Figs. 2 and S2). This provides an upper bound to the size of the region where a significant amount of WNT could be present, and thus constrains the diffusion length of WNT, $\ell_w = \sqrt{\eta_w/\delta_w}$ (Eq. (M2)). We use $\eta_w = 0.025\ \mu m^2/s$ which gives a diffusion length $\ell_w$ of approximately $31\ \mu m$ with $\delta_w = 1/11\ h^{-1}$. This translates into a dimensionless diffusivity $D_w = 0.001$. Further discussion of the impact of WNT diffusivity can be found in Sec. 4.3.

The pluripotent identity $P$ is a baseline state and does not play a central role to our cell fate network. Therefore we prescribe the parameters for its dynamics with the following considerations. On one hand, without preference to differentiate, cells should recover pluripotency in reasonable time to small perturbations and maintain weak repression of other fates. On the other hand, if any of the other fates is activated, there should be a strong inhibition on the pluripotent identity so cells differentiate. Hence, we assume

$$l_{xp} = l_{mp} = 0.8,\ l_{px} = l_{pm} = 0.2,\ r_{xp} = r_{mp} = r_{px} = r_{pm} = 8,\ k_p = 0.5,\ d_p = 1.$$

We expect that small variations of these parameters will not change the qualitative behaviour of the model.

#### 3.2 Fitting gradient profiles

Given the above parameters, we can calculate the static BMP4 response profile $B(y)$ and the evolution of the WNT profile $W(y,t)$ over time along the chamber length under any given BMP4 gradient $b(y)$ by Eqn. M7 and solving Eqn. M8. Then for each of the three different maximum BMP4 concentrations ($q = 2.5, 5, 10$) used in our experiments, we can simulate the evolution of cell fates $X,\ M,\ P$ and fit the $X,\ M$ profiles against experimental data (Fig. 2). This allows us to determine the values of related free parameters, which we assemble in the following vector

$$\mathbf{p} = (n_x, k_x, r_x, l_x, \bar{n}_x, \bar{k}_x, \bar{a}_x, d_x, n_m, k_m, r_m, l_m, d_m).$$

To carry out the fitting, we parse the experimental data first. Fractions of different cell types in the total population are extracted from fluorescence intensities in experimental images based on Table M1 (also see Fig.

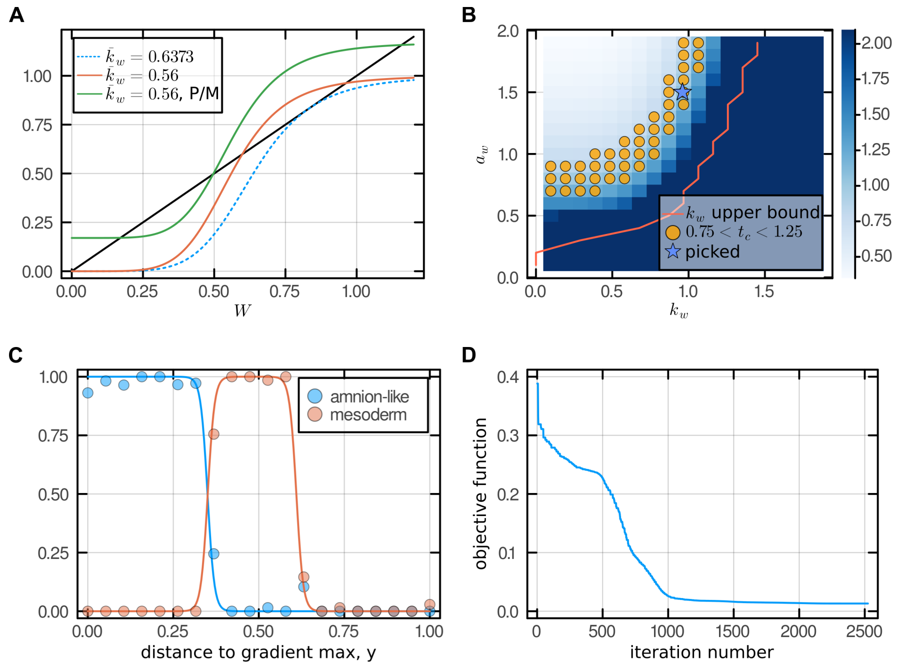


Figure M1: **Model parametrisation. A**. Prescribed bistability in WNT self-activation. Black line: WNT decay rate; blue dashed curve: WNT self activation rate at the critical $\bar{k}_w$ value for bistability; red curve: WNT self activation rate at the chosen $\bar{k}_w$ value when bistability is enabled; green curve: WNT production rate (including self activation and activation by BMP) with the chosen parameters ($\bar{n}_w,\ \bar{k}_w,\ n_w,\ k_w,\ a_w$) at the experimentally measured pluripotent/mesodermal fate boundary. **B**. Dimensionless duration $t_c$ (colour bar) of BMP4 stimulation after which WNT is self-sustaining in the $\mu$TransWell stimulation, as a function of WNT activation parameters. The red curve represents the maximum allowable $k_w$ that admits a sufficiently high $W$ which can self-sustain after stimulation is removed. Yellow markers are those with $0.75 < t_c < 0.25$. $b_s = 10,\ d_w = 1$. **C**. Reconstructed cell identity fractions from an example dataset of fluorescence intensities in the gradient chip. **D**. Convergence of the Nelder-Mead algorithm in parameter fitting. Preselected parameters, initial guesses and fitted values for other parameters are in Table M3.

2). Within the gradient, we seek the two types of fate transitions, namely from extra-embryonic to mesodermal and from mesodermal to pluripotent identities, in the sense of fate preferences at 50%. These transitions are found to always occur at $b_1 \approx 1.73$ and $b_2 \approx 1.00$, respectively (Fig. 2H). Then for each BMP4 maximum concentration $q_j,\ j = 1, 2, 3$, using the quadratic gradient profile, we can calculate the corresponding spatial locations $y_{i,j} = 1 - \sqrt{b_i/q_j},\ i = 1, 2$ of fate transition. To reduce measurement noise and obtain smooth fate profiles for fitting, since these boundaries are sharp, we reconstruct the target cell fate likelihoods as sigmoidal profiles in the following forms

$$\hat{X}_j(y) = \frac{1}{2}\left[1 - \tanh\left(\frac{y - y_{1,j}}{\epsilon}\right)\right], \tag{M15}$$

$$\hat{M}_j(y) = \frac{1}{2}\left[\tanh\left(\frac{y - y_{1,j}}{\epsilon}\right) - \tanh\left(\frac{y - y_{2,j}}{\epsilon}\right)\right], \tag{M16}$$

where $\epsilon = 0.02$ is a chosen parameter to control the width of the fate boundaries. We demonstrate an example of this reconstruction process compared to a specific experimental dataset in Fig. M1C, showing excellent agreement between the reconstructed (solid curves) and measured cell identities (markers).

With the preset parameters from Sec. 3.1 and the current free parameter set $\mathbf{p}$, for the $j$-th BMP4 gradient strength $q_j$, we can compute the likelihoods for extra-embryonic and mesodermal cells $X_j(y, t_g)$ and $M_j(y, t_g)$, respectively, at the given time of measurement ($t_g = 72\ h$) in the gradient chip experiments. The following objective (cost) function measures the difference of these profiles from the reconstructed experimental data:

$$f(\mathbf{p}) = \begin{cases} 10, & \text{if } k_x < 0, \bar{k}_x < 0, k_m < 0, l_x < 0, \text{ or } l_m < 0, \\ \displaystyle\sum_{j=1}^{3}\left[\int_0^1 \left(X_j(y,t_g) - \hat{X}_j(y)\right)^2 + \left(M_j(y,t_g) - \hat{M}_j(y)\right)^2 dy\right], & \text{otherwise.} \end{cases} \tag{M17}$$

Note that $f(\mathbf{p})$ imposes a penalty if any of the parameters $k_x, \bar{k}_x, k_m, l_x$, or $l_m$ violates the non-negativity constraint. Numerically, the integrals in Eqn. (M17) are approximated using the trapezoidal rule on the same

| | $q = 2.5$ | | $q = 5$ | | $q = 10$ | |
|---|---|---|---|---|---|---|
| experiment | 0.169 | 0.367 | 0.412 | 0.553 | 0.585 | 0.684 |
| simulation | 0.170 | 0.366 | 0.413 | 0.555 | 0.584 | 0.691 |

Table M2: Comparison of locations of cell fate transitions in the BMP gradient between experimental data and simulations. The values are normalised distances to the gradient max, $y$. Using parameters from Table. M3.

computational grid used to solve Eqn. (M8). Note that the cost function only considers the $X$ and $M$ fates, and excludes cells outside the stimulation window.

To minimise this objective function and evolve the estimated free parameters, we adopt the Nelder-Mead method, an iterative, derivative-free optimisation algorithm based on simplex reflections[19–22]. For a non-convex objective function, the algorithm allows for temporary increases in the objective value so as to escape local optima. We use an implementation of the Nelder-Mead method in the `Julia`[17] package `Optimization.jl`[20] and `Optim.jl`[23]. The energy landscape of $f(\mathbf{p})$ associated with the model is complex, and we identify a set of proper initial conditions by trial and error. Figure M1D shows the reduction in the objective function $f(\mathbf{p})$ against the iteration number as the optimisation algorithm progresses, and convergence is evidenced.

Finally, the evolution of the endoderm fate $E$ is slaved to that of the mesoderm $M$ fate and WNT. Therefore, once we obtain a set of free parameters $\mathbf{p}$ that fit the experimental data, we manually pick the parameters for Eqn. (M11). Again we expect the results will not change significantly with small variations in these parameters provided that they respect the qualitative behaviour of $E$. The fitted parameters, as well as the pre- and post-selected parameters, are summarised in Table M3.

### 4 Extended discussions of results

#### 4.1 Cell fates and their evolution

Using the preselected and fitted parameters in Table M3, we can reproduce quantitatively the cell identity profiles in the static BMP4 gradients as well as the fate choices in the $\mu$TransWells (Fig. 5). For the gradient chips, we extract the fate transition locations from the simulations and compare to the mean values measured in the experiments in Table M2. For each $q$ value, the first column gives the coordinate where the mesodermal and amnion-like fates transition, calculated as the average coordinate of either identity likelihood crosses the level of 0.5. The second column is the location of pluripotent/mesodermal transition and is the average of coordinates of $P = 0.5$ and $M = 0.5$.

In addition to the fate profiles in the gradient at 72 hours, we can track the evolution of all cell identity markers in time for $q = 5$ (Fig. M2). The model prediction for the mesodermal fate during experimental time (Fig. M2B) agrees well with the experimental data in Fig. S2C. The computed evolution of likelihood for the pluripotent state $P$ (Fig. M2D) is also comparable to experimental observation (Fig. S2D), with the caveat that this state is relatively unimportant in our model and its parameters are postulated.

For the $\mu$TransWell, our simulations could recapitulate the time-dependent differentiation of the mesodermal fate outside the stimulation window (Fig. 5D). On one hand, our fitting procedure has guaranteed that WNT becomes self-sustaining through bistability of self-activation after a duration around $t_0 = 11\ h$ of BMP4 stimulation. This ensures mesoderm differentiation after BMP4 stimulation stops. On the other hand, our model with the fitted parameters predicts that self-activation of the extra-embryonic fate $X$ is insufficient to enable bistability ($\bar{n}_x = 0.0002951,\ \bar{k}_x = 3.372,\ \bar{a}_x = 0.1030$) and therefore, the amnion-like identity vanishes once BMP4 stimulation stops, also in agreement with experimental observation.

#### 4.2 Time scale of WNT dynamics

The decay rate of WNT activity $\delta_w$ has not been measured directly. As described in Sec. 3.1, our experiments have suggested the characteristic time scale of WNT dynamics to be around 11 hours since it is the threshold duration of BMP4 stimulation to induce self-sustained WNT activity. This is in agreement with a recent study on the temporal effects of BMP stimulation on cell differentiation[1], but is longer by one order of magnitude than the time scale adopted in an earlier study on the BMP and WNT signalling cascade[13]. To further probe the impact of this parameter in our model, we accelerate WNT dynamics by increasing $d_w$ from 1 to 10, and perform again the grid search of $k_w$ and $a_w$ values in $\mu$TransWell simulations to recover the measured critical BMP4 stimulation time $0.75 < t_c < 1.25$ (see Sec. 3.1 for detailed procedure). With much faster WNT dynamics ($d_w = 10$), the region of compatible parameters has shrunk considerably and is almost collocated with the theoretical upper bound of $k_w$ to admit the bistable non-zero level of WNT in absence of BMP4 (Fig. M3). In this case, the parameter values of $k_w$ and $a_w$ will be highly sensitive to perturbations, resulting in poor

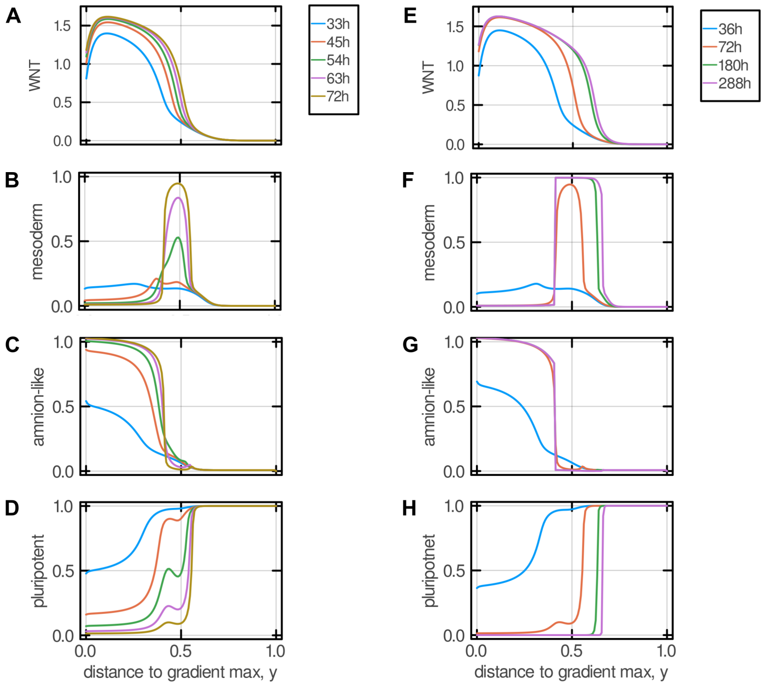


Figure M2: **Transient evolution of WNT and cell identity markers in a static BMP gradient in two time scales. A–D**. WNT and the likelihoods of different cells in experimental time scale up to 72 hours. **E–H**. Different species in a long time scale up to 288 hours. $q = 5$ and other parameters are from Table. M3. Time is presented in dimensional quantities for ease of comparison with experiments.

robustness of the model predictions. Therefore, we believe that a WNT decay time scale close to 11 hours is reasonable.

#### 4.3 Effects of WNT diffusion

Given that WNT is known to diffuse[11–13], we have prescribed a WNT diffusivity in our model, with a magnitude of $\eta_w = 0.025\ \mu m^2/s$ based on the reported average WNT diffusivity ($0.04\ \mu m^2/s$) in *Xenopus* embryos[18]. This small diffusion coefficient is attributed to the majority of WNT being bound to cell surfaces[18]. Consistently, our experimental observations in the $\mu$TransWell where the spread of the WNT reporter TCF/LEF:d2YFP is comparable to the spread of the stable BMP reporter BRE:YFP (Fig. 4G,H) argue for a limited extent of WNT diffusion in our microfluidic system. As a reference, our WNT diffusivity is still a few times larger than that assumed in Chhabra *et al.*[13].

To further examine the role of WNT diffusion, we carry out a sensitivity analysis by varying WNT diffusivity and keeping the other parameters constant as in Table M3. Increasing WNT diffusion rounds the fate boundaries and expands the width of the mesoderm bands in the middle of the BMP4 gradient and outside the

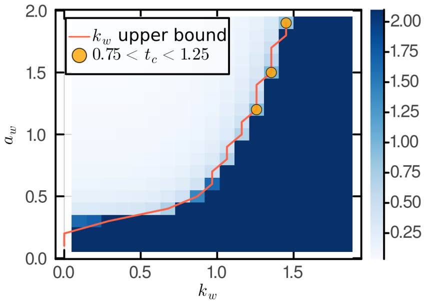


Figure M3: **WNT activation parameters with quick WNT relaxation.** Viable $a_w$ and $k_w$ values to fit $\mu$TransWell simulation threshold timing at $d_w = 10$. Other parameters: $b_s = 10,\ n_b = 1.355,\ k_b = 0.8233,\ n_w = 4,\ \bar{n}_w = 6,\ \bar{k}_w = 0.56$.

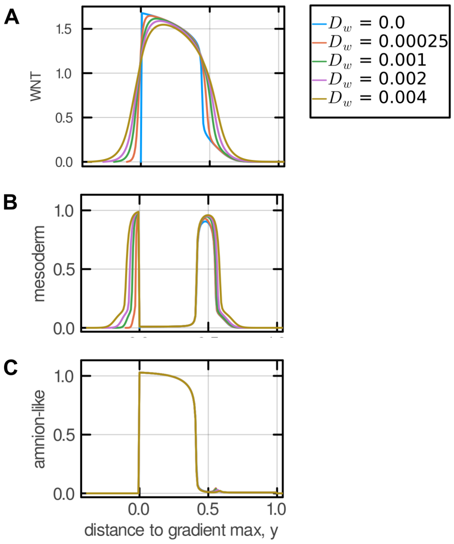


Figure M4: **Sensitivity to WNT diffusion.** Profiles of WNT (**A**), mesodermal $M$ (**B**) and extra-embryonic, amnion-like $X$ (**C**) identities in the static BMP gradient with $q = 5$ at 72 hours, with different WNT diffusivities. Other parameters are from Table M3.

gradient maximum, both of which agree with expectation (Fig .M4). In particular, the mesoderm size outside the static BMP gradient scales approximately with $\sqrt{D_w}$ (Fig. M4A, B), as expected because this region is entirely controlled by WNT diffusion. Other than these quantitative effects, the shape and evolution of the fate bands are not qualitatively altered (Fig. M4). In the extreme case of WNT diffusion being removed, i.e., setting $D_w = 0$, we have re-fitted the parameters to examine if we could still recover the experimental observations (fitting results reported in Table M4). Excellent agreement has been achieved, including the cell fate profiles at 72 hours in the BMP gradient chip (Fig. M5A) and their evolution over time (Fig. M5C, E), as well as cell identities in the $\mu$TransWell device (Fig. M5F–I).

Combining experimental and numerical evidence, we conclude that WNT diffusion is not a necessary condition and has limited impact in reproducing the experimental observations. However, the existence of a small WNT diffusion leads to a long-term shift of the pluripotent/mesodermal boundary, as detailed next in Sec 4.4.

#### 4.4 Transient cell fates in static BMP gradients

One question related to WNT diffusion is whether the cell fate markers in the BMP4 gradient actually reached steady state at 72 hours when the experiments were concluded. Because of the presence of a small WNT diffusivity, our model predicts that the evolution of cell fates within the gradient undergoes two distinct stages. The first is dominated by production and inhibition (reaction terms in Eqns. M8–M10), while diffusion plays a secondary role. The end of this stage largely sees the establishment of the observed fate bands already (Fig. M2A–D). Model predictions are in close agreement with experiments (Fig. S2). During the second stage, diffusion drives a slow invasion of the WNT front from the high-BMP end towards the low-BMP end, thus causing the migration of the pluripotent/mesodermal fate boundary towards the gradient minimum. This WNT front migration in the second stage is well-known in reaction-diffusion systems[24]. The slow motion of the mesoderm front, however, is only observable during much longer time after the amnion-like and mesoderm layers form, as clearly demonstrated in Fig. M2E–H. Practically, it is challenging to be identified in experiments due to the requirement of maintaining stable cell conditions for a much longer duration than this developmental stage in hESCs.

With the parameter set for Figs. 5 and M2 (Table M3) and $q = 5$, the model predicts that in the BMP gradient the WNT front, and thus the pluripotent/mesoderm fate boundary, eventually stops at a distance of $\Delta y \approx 0.1$ from the experimentally measured fate transition towards the gradient minimum. Note that the additional displacement is comparable to the mesoderm band in size, and is hence significant. This equilibrium position lies where the speed of the WNT wave front vanishes and depends on the choices of $n_w, a_w,\ k_w,\ \bar{n}_w,\ \bar{k}_w$ given the BMP stimulation response ($B$). The five parameters are preselected following the procedure in Sec. 3.2. In fact, we could alternatively adopt a different set of parameters that place the equilibrium WNT position at a different location. As an example, in Table M5 we present a set of WNT activation and self-activation parameters such that the WNT front will migrate beyond the BMP gradient minimum. In consequence, in long

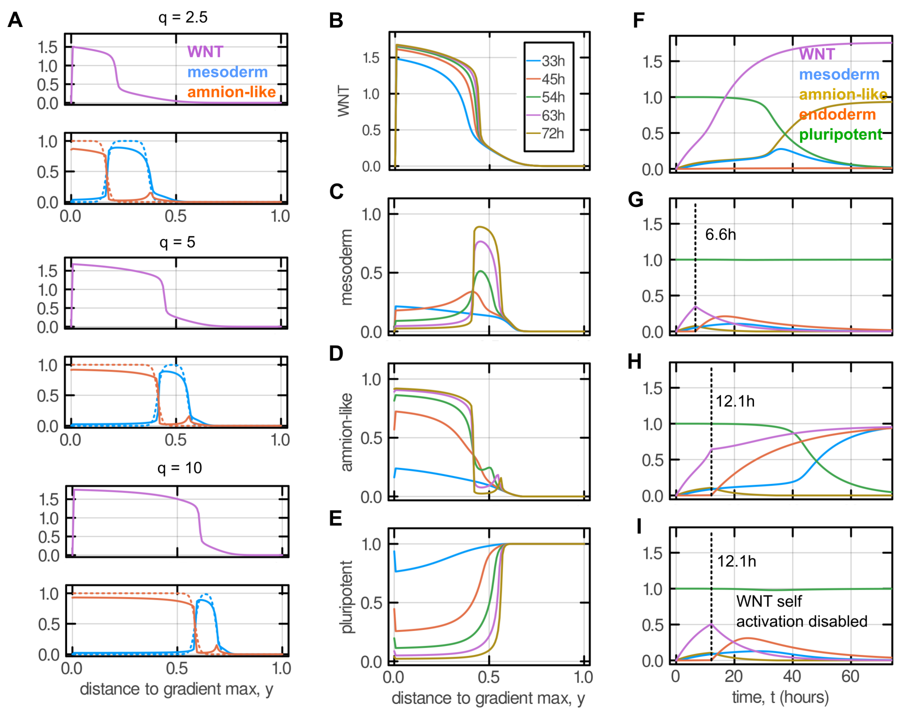


Figure M5: **Simulation results obtained from re-fitted parameters without WNT diffusion** ($D_w = 0$). **A**. Profiles of WNT activity and the mesodermal and amnion-like fate likelihoods in the static BMP4 gradient at different gradient maximum values ($q = 2.5,\ 5,\ 10$). The dashed curves are target fate bands extracted from the experiments, as explained in Sec. 3.2. **B–E**. Time evolution of WNT activity and the likelihoods for mesodermal, amnion-like and pluripotent cells in the BMP4 gradient with $q = 5$. **F–I**. Evolution of WNT activity and likelihoods for all cell types in the $\mu$TransWell under different durations of BMP4 stimulation. $b_s = 10$. The conditions are the same as in Fig. 5D. **F**. Continuous stimulation. **G–I**. The dashed line marks when stimulation stops. **I**. WNT self-activation is disabled. Preselected and fitted parameters are in Table M4. Time is presented in dimensional quantities for ease of comparison with experiments.

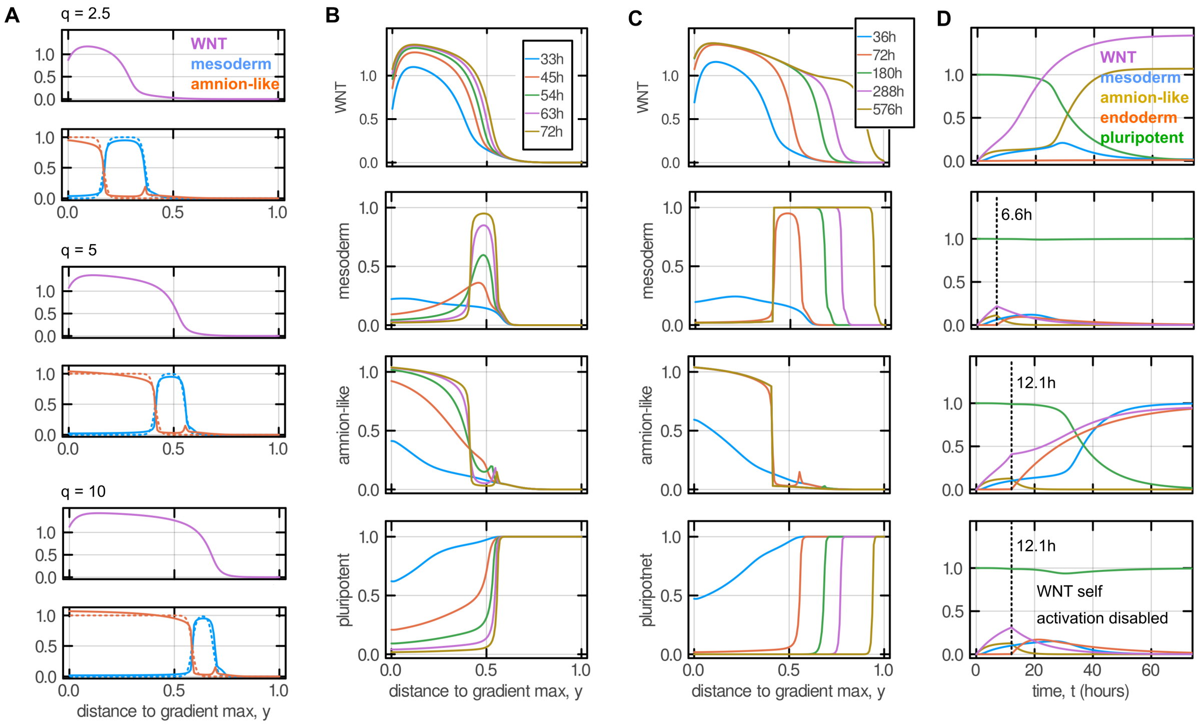


Figure M6: **Simulation results with WNT steady-state front location outside the stimulation window of the BMP gradient.** **A**. Profiles of WNT activity and the mesodermal and amnion-like fate likelihoods in the static BMP4 gradient at different gradient maximum values ($q = 2.5,\ 5,\ 10$). The dashed curves are target fate bands extracted from the experiments, as explained in Sec. 3.2. **B**. Evolution of WNT activity and cell identity profiles in experimental time up to 72 hours in the static BMP4 gradient with $q = 5$. **C**. Evolution of WNT activity and cell type likelihoods in a longer time scale. **D**. Evolution of WNT activity and cell type likelihoods in the $\mu$TransWell under different durations of BMP4 stimulation (continuous, stopping at 6.6 hours and 12.1 hours). Preselected and fitted parameters are in Table M5. Time is presented in dimensional quantities for ease of comparison with experiments.

time the pluripotent/mesoderm identity boundary will sweep through the entire BMP stimulation window (Fig. M6C). In this case, we could still fit the rest of the parameters and obtain model predictions in good agreement with the experimental results. These again include the cell state profiles in the gradient at 72 hours (Fig. M6A), and their evolution in both the gradient (Fig. M6B) and the $\mu$TransWells (Fig. M6D).

In either case of long-term WNT front behaviour, to ensure that the model output matches experimental observation, the equilibrium WNT front location must lie towards the side of gradient minimum relative to the pluripotent/mesoderm boundary. It follows that at the target pluripotent/mesoderm boundary in the BMP4 gradient, the velocity of the WNT front transitioning from the low to high levels points at the gradient minimum (green solid curve in Fig. M1A). This condition needs to be satisfied when choosing the WNT activation and self activation parameters $n_w, a_w,\ k_w,\ \bar{n}_w,\ \bar{k}_w$, which has been taken into account in our parametrisation procedure (see Sec. 3.1).

Whether the WNT front arrives at equilibrium inside the stimulation window or moves outside, the second stage takes a longer time scale than experimental time. This is suggested in the estimate that the front speed is on the order of $\sqrt{\eta_w \delta_w} = 0.0029\ mm/h$ by dimensional analysis. On one hand, it is constrained by the critical duration for WNT self-sustainability measured from the $\mu$TransWells. On the other hand, as the WNT front approaches steady state, its speed vanishes. To validate this prediction, we fit again the parameters in $\mathbf{p}$ by minimising the difference between measured cell fate profiles and simulated ones at *steady state*, instead of at 72 hours after stimulation starts. The resulting parameter values are summarised in Table M6. Fig. M7A shows the steady-state species which follow the target profiles well. However, it takes one order of magnitude longer time than experimental to reach equilibrium, while at 72 hours, the likelihood of the mesodermal fate $M$ is below 0.5, not considered differentiated (Fig. M7B, in comparison to Figs. S2 and M2C).

In conclusion, our model, constrained by the experimental data, shows that the measured cell fate profiles are transient in nature. The pluripotent/mesoderm boundary, together with the underlying wave front of the morphogen WNT, continues moving slowly. The fronts are unlikely to reach steady state within experimental time given the small WNT diffusivity and long relaxation time. In this sense, although the differentiation of hESCs induced in our BMP gradient displays a French flag pattern, the underlying system, driven by the

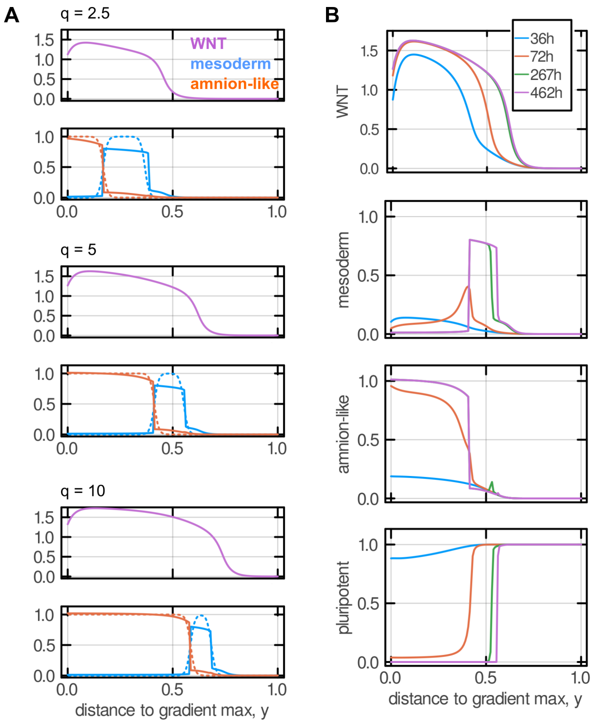


Figure M7: **Simulation results with parameters fitted at steady state in the BMP gradient. A**. Profiles of WNT activity and the mesodermal and amnion-like fate likelihoods in the static BMP4 gradient at different gradient maximum values ($q = 2.5,\ 5,\ 10$). The dashed curves are target fate bands extracted from the experiments, as explained in Sec. 3.2. **B**. Evolution of WNT activity and cell identity profiles in time up to steady state. Preselected and fitted parameters are in Table M6. Time is presented in dimensional quantities for ease of comparison with experiments.

secondary morphogen WNT, is different from the classical French flag picture[25].

#### 4.5 Interaction of cell fate determinants

The extra-embryonic $X$ and mesodermal $M$ cell identities, together with the morphogens BMP4 and WNT, form the core interactions in our cell fate network (Fig. 5B). We have also considered the pluripotent state $P$ as a baseline identity, whose cross repression with $M$ and $X$ contribute to the formation of sharp fate boundaries. The inclusion of the endoderm fate $E$ is for completeness. Because we neglect its mutual inhibition with other fates, it becomes slaved to WNT and BMP4. We use prescribed parameters for $P$ and $E$ as these are not essential ingredients while fitting the parameters of the core components involving $M$ and $X$ (Sec. 3). We have adopted a simplified dynamics for the morphogen WNT (Sec. 1).

Notice that in the fitted parameters (Tables M3–M4, M6), the Hill coefficient $n_x$ for extra-embryonic fate activation by BMP4 is often high, suggesting strong cooperativity. Biologically this is likely due to detailed signalling and feedback loops that are not resolved in our simplified cell fate network (Fig. 5B). The high $n_x$ value contributes to the sharp fate boundaries of amnion-like tissue. This is possibly further refined by mutual inhibition between the extra-embryonic fate $X$ and other identities, particularly the mesodermal fate $M$ and the pluripotent state $P$. The details of such fate interactions could be worth probing in future studies.

In the model, we have included self activation of the amnion-like identity $X$ in the form of a Hill function with coefficients $(\bar{n}_x, \bar{k}_x, \bar{a}_x)$ (Eqn. M9). Fitting in several different setups against our experimental data has consistently resulted in small (Tables M3, M5–M6), or even negative (Table M4) $\bar{a}_x$ values, suggesting negligible or insignificant effects of extra-embryonic cell self-activation. This is in agreement with our $\mu$TransWells experiments in that cells not receiving continuous BMP stimulation cannot consistently differentiate into the amnion-like fate. Our model indicates that the amnion-like identity is only established after a duration close to 30 hours of BMP4 induction in the $\mu$TransWell (Figs. 5D, M5F, M6D). If stopping BMP at this time, due to the lack of bistability, eventually the extra-embryonic likelihood $X$ will be lost but its transient presence may be able to maintain the identity long enough for later developmental stage. This result also matches the earlier study on hESC differentiation by BMP4 in regular tissue culture dishes by Camacho-Aguilar *et al.* who have found that the coefficient for the self-activation Hill function for CDX2-marked cells is 0.0922 after fitting to experiments[1]. Indeed, in their study, consistent extra-embryonic differentiation was only observed after 30 hours of BMP4 stimulation and observation stopped at 48 hours. More detailed characterisation of signalling and differentiation related to the amnion-like fate could be useful to compare with our conclusion.

### 5 Summary

With a combination of experiments and numerical simulations, we show that our model of a simplified cell fate network (Fig. 5) captures the patterning logic of the hESCs. Our experimental measurements provide strong constraints on WNT bistability and dynamics. This allows us to parametrise the model by selecting a number of key values and fitting the rest against experimental data. The model quantitatively recapitulates cell differentiation when exposed to both graded and uniform BMP stimulations. In the BMP gradient experiment, the model demonstrates a classical French flag pattern featuring three bands of different cell identities inside the stimulation window, albeit driven by the secondary morphogen WNT and cell fate interactions. Our model further demonstrates that long WNT dynamics with weak diffusion leads to a slow WNT bistable front motion in the gradient experiment and prevents the establishment of a fully steady pattern during the few days of the experiment, distinct from the classical French flag theory.

Future detailed investigations, particularly measurement of WNT and NODAL dynamics in the early development of human embryos and characterisation of extra-embryonic autoregulation, will be required to fully clarify the interplay of WNT time scales and those of cell differentiation. The wave front propagation of WNT signalling and the pluripotent/mesodermal fate boundary is reminiscent of the WNT and NODAL waves travelling at constant speeds across circular hESC colonies under BMP stimulation[13]. In that case, BMP treatment was presumably concentrated at the colony edge due to inaccessibility to the basal receptors, and thus resembled more a step stimulation. The waves were considered to be caused by reaction-diffusion outside the Turing instability regime, but closer examinations were desirable. A recent study probed the differentiation of hESCs and identified gastrulation-like events including primitive streak formation when the epiblast was placed along side an amnion-like tissue[10]. It highlights that BMP signalling could in turn originate from extra-embryonic tissues and NODAL signalling is essential in cell differentiation, suggesting more sophisticated signalling and cell fate interactions.

With the large number of parameters, unsurprisingly there is *sloppiness* in their values in our model[26]. We have showcased three sets of parameters (Tables M3–M5) whose simulation results all recover the observed cell fate marker distribution in the BMP gradient and the $\mu$TransWell cell identity evolution. However, we have also demonstrated that the main effect of the small WNT diffusivity to drive a long-time fate boundary migration is robust to parameter variations. Hence, we expect that our qualitative predictions to hold despite the uncertainty in individual parameters[26]. Taken together, combining microfluidic devices capable of applying well-controlled temporal and spatial morphogen stimulation with mechanistic models offers a powerful tool to decipher tissue patterning.

##### Data availability

`Julia` codes for numerical simulation, fitting and data analysis are available in https://github.com/bsorre/wyatt_gradients.

**Measured parameters**

| Parameter | Value | Physical interpretation | Nondimensionalisation |
|---|---|---|---|
| $n_b$ | 1.355 | BMP4 response Hill coefficient | |
| $k_b$ | 0.8233 | BMP4 response Hill dissociation constant | $k_b = K_b/b_0$ |

**Preselected parameters for WNT dynamics**

| Parameter | Value | Physical interpretation | Nondimensionalisation |
|---|---|---|---|
| $D_w$ | 0.001 | $W$ diffusivity | $D_w = \eta_w/(\delta_w L_0^2)$ |
| $n_w$ | 4 | $W$ activation Hill coefficient | |
| $k_w$ | 0.96 | $W$ activation Hill dissociation constant | $k_w = K_w/A_b$ |
| $a_w$ | 1.5 | $W$ activation strength | $a_w = A_w/\bar{A}_w$ |
| $\bar{n}_w$ | 6 | $W$ self-activation Hill coefficient | |
| $d_w$ | 1 | $W$ decay rate | $d_w = \delta_w t_0$ |
| $\bar{k}_w$ | 0.56 | $W$ self-activation Hill dissociation constant | $\bar{k}_w = \delta_w \bar{K}_w/\bar{A}_w$ |

**Preselected parameters related to pluripotent state $P$**

| Parameter | Value | Physical interpretation | Nondimensionalisation |
|---|---|---|---|
| $n_p$ | 1 | $P$ self-activation Hill coefficient | |
| $k_p$ | 0.5 | $P$ self-activation Hill dissociation constant | $k_p = \delta_p K_p/A_p$ |
| $r_{xp}$ | 8 | inhibition Hill coefficient of $X$ by $P$ | |
| $l_{xp}$ | 0.8 | baseline inhibition factor of $X$ by $P$ | $l_{xp} = \delta_p L_{xp}/A_p$ |
| $r_{mp}$ | 8 | inhibition Hill coefficient of $M$ by $P$ | |
| $l_{mp}$ | 0.8 | baseline inhibition factor of $M$ by $P$ | $l_{mp} = \delta_p L_{mp}/A_p$ |
| $r_{px}$ | 8 | inhibition Hill coefficient of $P$ by $X$ | |
| $l_{px}$ | 0.2 | baseline inhibition factor of $P$ by $X$ | $l_{px} = \delta_x L_{px}/A_x$ |
| $r_{pm}$ | 8 | inhibition Hill coefficient of $P$ by $M$ | |
| $l_{pm}$ | 0.2 | baseline inhibition factor of $P$ by $M$ | $l_{pm} = \delta_m L_{pm}/A_m$ |
| $d_p$ | 1 | $P$ decay rate | $d_p = \delta_p t_0$ |

**Fitted parameters for amnion-like $X$ and mesoderm $M$ fates**

| Parameter | Initial value | Fitted value | Physical interpretation | Nondimensionalisation |
|---|---|---|---|---|
| $n_x$ | 4.0 | 11.40 | $X$ activation Hill coefficient | |
| $k_x$ | 0.5 | 0.6589 | $X$ activation Hill dissociation constant | $k_x = K_x/A_b$ |
| $r_x$ | 5.5 | 3.233 | inhibition Hill coefficient of $X$ by $M$ | |
| $l_x$ | 2.1 | 0.2421 | baseline inhibition factor of $X$ by $M$ | $l_x = \delta_m L_x/A_m$ |
| $\bar{n}_x$ | 1.6 | 0.0002951 | $X$ self-activation Hill coefficient | |
| $\bar{k}_x$ | 1.5 | 3.372 | $X$ self-activation Hill dissociation constant | $\bar{k}_x = \delta_x \bar{K}_x/A_x$ |
| $\bar{a}_x$ | 0.2 | 0.1030 | $X$ self-activation production rate | $\bar{a}_x = \bar{A}_x/A_x$ |
| $d_x$ | 2.4 | 1.756 | $X$ decay rate | $d_x = \delta_x t_0$ |
| $n_m$ | 4.0 | 2.427 | $M$ activation Hill coefficient | |
| $k_m$ | 0.2 | 0.08066 | $M$ activation Hill dissociation constant | $k_m = \delta_w K_m/\bar{A}_w$ |
| $r_m$ | 2.9 | 2.967 | inhibition Hill coefficient of $M$ by $X$ | |
| $l_m$ | 0.2 | 0.2087 | baseline inhibition factor of $M$ by $X$ | $l_m = \delta_x L_m/A_x$ |
| $d_m$ | 0.61 | 1.455 | $M$ decay rate | $d_m = \delta_m t_0$ |

**Postulated parameters for endoderm fate $E$**

| Parameter | Value | Physical interpretation | Nondimensionalisation |
|---|---|---|---|
| $n_e$ | 4 | $E$ activation Hill coefficient | |
| $k_e$ | 0.2 | $E$ activation Hill dissociation constant | $k_e = \delta_w K_e/\bar{A}_w$ |
| $r_e$ | 2 | inhibition Hill coefficient of $E$ by $B$ | |
| $l_e$ | 0.1 | baseline inhibition factor of $E$ by $B$ | $l_e = L_e/A_b$ |
| $d_e$ | 0.5 | $E$ decay rate | $d_e = \delta_e t_0$ |

Table M3: Preselected and fitted model parameters. See the main text and Figs. 5 and M2 for simulation results.

**Preselected parameters for WNT dynamics**

| Parameter | Value | Physical interpretation | Nondimensionalisation |
|---|---|---|---|
| $D_w$ | 0.001 | $W$ diffusivity | $D_w = \eta_w/(\delta_w L_0^2)$ |

**Fitted parameters for amnion-like $X$ and mesoderm $M$ fates**

| Parameter | Initial value | Fitted value | Physical interpretation | Nondimensionalisation |
|---|---|---|---|---|
| $n_x$ | 4.0 | 6.871 | $X$ activation Hill coefficient | |
| $k_x$ | 0.5 | 0.5749 | $X$ activation Hill dissociation constant | $k_x = K_x/A_b$ |
| $r_x$ | 5.5 | 7.544 | inhibition Hill coefficient of $X$ by $M$ | |
| $l_x$ | 2.1 | 0.3598 | baseline inhibition factor of $X$ by $M$ | $l_x = \delta_m L_x/A_m$ |
| $\bar{n}_x$ | 1.6 | 1.536 | $X$ self-activation Hill coefficient | |
| $\bar{k}_x$ | 1.5 | 1.273 | $X$ self-activation Hill dissociation constant | $\bar{k}_x = \delta_x \bar{K}_x/A_x$ |
| $\bar{a}_x$ | 0.2 | -0.07226 | $X$ self-activation production rate | $\bar{a}_x = \bar{A}_x/A_x$ |
| $d_x$ | 2.4 | 1.435 | $X$ decay rate | $d_x = \delta_x t_0$ |
| $n_m$ | 4.0 | 3.414 | $M$ activation Hill coefficient | |
| $k_m$ | 0.2 | 0.07743 | $M$ activation Hill dissociation constant | $k_m = \delta_w K_m/\bar{A}_w$ |
| $r_m$ | 2.9 | 5.141 | inhibition Hill coefficient of $M$ by $X$ | |
| $l_m$ | 0.2 | 0.3514 | baseline inhibition factor of $M$ by $X$ | $l_m = \delta_x L_m/A_x$ |
| $d_m$ | 0.61 | 0.9476 | $M$ decay rate | $d_m = \delta_m t_0$ |

Table M4: Fitted parameters for the model with no WNT diffusion ($D_w = 0$). Refer to Table M3 for some omitted parameters that are not changed. See Fig. M5 for simulation results.

**Preselected parameters for WNT dynamics**

| Parameter | Value | Physical interpretation | Nondimensionalisation |
|---|---|---|---|
| $D_w$ | 0.001 | $W$ diffusivity | $D_w = \eta_w/(\delta_w L_0^2)$ |
| $n_w$ | 4 | $W$ activation Hill coefficient | |
| $k_w$ | 1 | $W$ activation Hill dissociation constant | $k_w = K_w/A_b$ |
| $a_w$ | 1 | $W$ activation strength | $a_w = A_w/\bar{A}_w$ |
| $\bar{n}_w$ | 4 | $W$ self-activation Hill coefficient | |
| $d_w$ | 1 | $W$ decay rate | $d_w = \delta_w t_0$ |
| $\bar{k}_w$ | 0.4274 | $W$ self-activation Hill dissociation constant | $\bar{k}_w = \delta_w \bar{K}_w/\bar{A}_w$ |

**Fitted parameters for amnion-like $X$ and mesoderm $M$ fates**

| Parameter | Initial value | Fitted value | Physical interpretation | Nondimensionalisation |
|---|---|---|---|---|
| $n_x$ | 4.0 | 3.157 | $X$ activation Hill coefficient | |
| $k_x$ | 0.5 | 0.6719 | $X$ activation Hill dissociation constant | $k_x = K_x/A_b$ |
| $r_x$ | 5.5 | 2.391 | inhibition Hill coefficient of $X$ by $M$ | |
| $l_x$ | 2.1 | 0.2950 | baseline inhibition factor of $X$ by $M$ | $l_x = \delta_m L_x/A_m$ |
| $\bar{n}_x$ | 1.6 | 1.809 | $X$ self-activation Hill coefficient | |
| $\bar{k}_x$ | 1.5 | 0.1264 | $X$ self-activation Hill dissociation constant | $\bar{k}_x = \delta_x \bar{K}_x/A_x$ |
| $\bar{a}_x$ | 0.2 | 0.3177 | $X$ self-activation production rate | $\bar{a}_x = \bar{A}_x/A_x$ |
| $d_x$ | 2.4 | 4.047 | $X$ decay rate | $d_x = \delta_x t_0$ |
| $n_m$ | 4.0 | 5.502 | $M$ activation Hill coefficient | |
| $k_m$ | 0.2 | 0.06741 | $M$ activation Hill dissociation constant | $k_m = \delta_w K_m/\bar{A}_w$ |
| $r_m$ | 2.9 | 3.047 | inhibition Hill coefficient of $M$ by $X$ | |
| $l_m$ | 0.2 | 0.2833 | baseline inhibition factor of $M$ by $X$ | $l_m = \delta_x L_m/A_x$ |
| $d_m$ | 0.61 | 1.383 | $M$ decay rate | $d_m = \delta_m t_0$ |

Table M5: Alternative set of preselected and fitted model parameters that allow the WNT front and the pluripotent/mesoderm boundary to sweep through the entire stimulation window in the BMP gradient in the long term. Refer to Table M3 for some omitted parameters that are not changed. See Fig. M6 for results.

**Fitted parameters for amnion-like $X$ and mesoderm $M$ fates**

| Parameter | Initial value | Fitted value | Physical interpretation | Nondimensionalisation |
|---|---|---|---|---|
| $n_x$ | 4.0 | 10.36 | $X$ activation Hill coefficient | |
| $k_x$ | 0.5 | 0.6306 | $X$ activation Hill dissociation constant | $k_x = K_x/A_b$ |
| $r_x$ | 5.5 | 3.467 | inhibition Hill coefficient of $X$ by $M$ | |
| $l_x$ | 2.1 | 0.4286 | baseline inhibition factor of $X$ by $M$ | $l_x = \delta_m L_x/A_m$ |
| $\bar{n}_x$ | 1.6 | 1.229 | $X$ self-activation Hill coefficient | |
| $\bar{k}_x$ | 1.5 | 0.06469 | $X$ self-activation Hill dissociation constant | $\bar{k}_x = \delta_x \bar{K}_x/A_x$ |
| $\bar{a}_x$ | 0.2 | 0.03262 | $X$ self-activation production rate | $\bar{a}_x = \bar{A}_x/A_x$ |
| $d_x$ | 2.4 | 1.048 | $X$ decay rate | $d_x = \delta_x t_0$ |
| $n_m$ | 4.0 | 2.581 | $M$ activation Hill coefficient | |
| $k_m$ | 0.2 | 0.5630 | $M$ activation Hill dissociation constant | $k_m = \delta_w K_m/\bar{A}_w$ |
| $r_m$ | 2.9 | 5.096 | inhibition Hill coefficient of $M$ by $X$ | |
| $l_m$ | 0.2 | 0.4466 | baseline inhibition factor of $M$ by $X$ | $l_m = \delta_x L_m/A_x$ |
| $d_m$ | 0.61 | 1.079 | $M$ decay rate | $d_m = \delta_m t_0$ |

Table M6: Fitted model parameters when assuming the cell identities in the BMP gradient have reached steady state. Refer to Table M3 for some omitted parameters that are not changed. See Fig. M7 for results.
